# *Plasmodium vivax* Malaria viewed through the lens of an eradicated European strain

**DOI:** 10.1101/736702

**Authors:** Lucy van Dorp, Pere Gelabert, Adrien Rieux, Marc de Manuel, Toni de-Dios, Shyam Gopalakrishnan, Christian Carøe, Marcela Sandoval-Velasco, Rosa Fregel, Iñigo Olalde, Raül Escosa, Carles Aranda, Silvie Huijben, Ivo Mueller, Tomàs Marquès-Bonet, François Balloux, M. Thomas P Gilbert, Carles Lalueza-Fox

## Abstract

The protozoan *Plasmodium vivax* is responsible for 42% of all cases of malaria outside Africa. The parasite is currently largely restricted to tropical and subtropical latitudes in Asia, Oceania and the Americas. Though, it was historically present in most of Europe before being finally eradicated during the second half of the 20th century. The lack of genomic information on the extinct European lineage has prevented a clear understanding of historical population structuring and past migrations of *P. vivax*. We used medical microscope slides prepared in 1944 from malaria-affected patients from the Ebro Delta in Spain, one of the last footholds of malaria in Europe, to generate a genome of a European *P. vivax* strain. Population genetics and phylogenetic analyses placed this strain basal to a cluster including samples from the Americas. This genome allowed us to calibrate a genomic mutation rate for *P. vivax*, and to estimate the mean age of the last common ancestor between European and American strains to the 15th century. This date points to an introduction of the parasite during the European colonisation of the Americas. In addition, we found that some known variants for resistance to anti-malarial drugs, including Chloroquine and Sulfadoxine, were already present in this European strain, predating their use. Our results shed light on the evolution of an important human pathogen and illustrate the value of antique medical collections as a resource for retrieving genomic information on pathogens from the past.

## Introduction

Malaria is a leading cause of infectious disease, responsible for an estimated 200 million infections annually, and around 429,000 fatal cases (World Health Organisation 2017). The disease is caused by several species of parasitic protozoans from the genus *Plasmodium*, which are transmitted by various species of mosquitoes from the genus *Anopheles*. Two species in particular - *P. falciparum* and *P. vivax* - are responsible for the majority of human infections worldwide. Although *P. falciparum* causes 99% of malaria deaths globally, *P. vivax* is the aetiological agent of 42% of all cases outside of Africa (Gething et al. 2011; World Health Organisation 2017).

Today, the endemicity of genus *Plasmodium* is restricted to tropical and subtropical latitudes, spanning large regions of East and South-East Asia, Sub-Saharan Africa, Central and South America and Melanesia (Battle et al. 2012; Howes et al. 2016; Battle et al. 2019; Weiss et al. 2019). However, malaria was historically present in most of Europe, from the Mediterranean to the southern shores of the Baltic Sea, and from southern Britain to European Russia (Huldén et al. 2005). Malaria was eradicated from all European countries during the second half of the 20th century (Hay et al. 2004), with Spain being one of its last footholds from which it was only declared officially eradicated in 1964 (Pletsch 1965). Nevertheless, even though *Plasmodium* is currently largely absent from Europe, its potential re-emergence has been identified as a plausible consequence of climate change (Petersen et al. 2013; Zhao et al. 2016).

Whilst historically being described as the “benign” form of malaria, *P. vivax* is increasingly recognized as a significant cause of disease and mortality (Tjitra et al. 2008; Price et al. 2009; Lacerda et al. 2012; Baird 2013). In stark contrast to *P. falciparum*, *P. vivax* is capable of producing recurrent malaria episodes from a single infection due to its resistant latent forms known as hypnozoites (Gonzalez-Ceron et al. 2013; Adekunle et al. 2015). This capacity also allows *P. vivax* to maintain itself in temperate climates, resting in a dormant state in the cold months when anopheline populations are in diapause, and creating a persistent presence of parasite reservoirs which can facilitate wide-spread transmission and recurrent long-term infections (Krotoski 1985; Gething et al. 2011; White 2011). Low parasite densities of *P. vivax* in mixed infections (Mayxay et al. 2004; Moreira et al. 2015) and the suggested high proportion of hypozonoite-derived clinical incidence (Price et al. 2009; Howes et al. 2016) means the global prevalence of *P. vivax* is likely systematically underestimated in comparison to the more well studied *P. falciparum*.

*P. vivax* is widely considered to have emerged in Sub-Saharan Africa, a region in which it is now at low prevalence (Liu et al. 2014; Gunalan et al. 2018; Twohig et al. 2019). From there, it is thought to have spread globally through a complex pattern of migration events by hitchhiking with its human host, as humans moved out of Africa (Culleton et al. 2011; Hupalo et al. 2016). The analysis of a geographically diverse dataset of 941 *P. vivax* mitochondrial DNA (mtDNA) genomes detected genetic links between strains from the Americas to those from Africa and South Asia, although potential contributions from Melanesia into the Americas were also identified (Rodrigues et al. 2018). However, the role of the worldwide colonial expansion of European countries in the global dispersal of *P. vivax* remains largely unknown, mainly due to the lack of available sequenced nuclear genomes from now-extinct European strains.

The recent discovery of a set of historic microscope slides with bloodstains from malaria-affected patients provides major opportunities to shed light on the evolution of *P. vivax*. The slides were prepared between 1942-1944 in the Ebro Delta (Spain), an area where the disease was transmitted by the mosquito species *Anopheles atroparvus*, a member of the *A. maculipennis* complex - still very common in the region - and provided the first retrieval of genetic material from historical European *P. vivax* (Gelabert et al. 2016) as well as a partial *P. falciparum* genome (de-Dios et al. 2019 *in press*). The complete mtDNA genome of this sample showed a close genetic affinity to the most common strains of present-day South and Central America, suggesting their introduction into the Americas was linked to Spanish colonial-driven transmission of European strains. However, mtDNA is a maternally inherited single locus and, in comparison to the entire genome, has limited power to reconstruct complex evolutionary histories.

In this paper, we extend these previous findings, by reporting the complete genome of an extinct European *P. vivax* obtained from the historical Ebro Delta microscope slides. Together with the recent publication of a more accurately annotated *P. vivax* reference genome (PvP01) (Auburn et al. 2016), this European genome provides new opportunities to resolve the historical dispersals of this parasite using genome-wide data. The availability of this complete genome also allows direct estimation of an evolutionary rate for *P. vivax*, making it possible to date the clustering of the historic European strain with modern global strains. Furthermore, it enables us to ascertain the presence of some resistance-alleles prior to the introduction of most anti-malaria drugs. This information is critical for further investigations into the evolution of the parasite, as well as predicting future emergences of drug-resistance mutations.

## Results

We generated shotgun Illumina sequence data from four archival blood slides derived from malaria patients sampled between 1942 and 1944 in Spain’s Ebro Delta. The majority of reads that mapped to *P. vivax* (89.09%) derived from a single slide dated to 1944. In total, 481,245 DNA reads (0.44% of the total reads generated) mapped to *P. vivax*, yielding a composite genome, we term Ebro-1944, at 1.4x coverage (1.28x deriving from the newly analysed 2017 slide), and spanning 66.42% of the (PvP01) reference. The mtDNA genome was recovered at 32x coverage (Supplementary fig. 1-3 and Supplementary Tables 1-2). Although working with low coverage ancient genomes is challenging, the fact that our genome is haploid and displays low levels of post-mortem sequencing errors suggests it can be reliably used in most evolutionary analyses.

To explore the phylogeographic affinities of the eradicated European strain we performed several population genomics analyses (Supplementary fig. 4-5 and Supplementary Table 3). A principal component analysis (PCA) applied to a global dataset of *P. vivax* showed strong geographic structure, with clusters separating 1) South East Asian/East Asian strains; 2) Oceanian strains (those from Malaysia were placed between the two clusters); 3) Indian and Madagascan strains and 4) those sampled from Central/South America (fig. 1a,b). The European sample (labelled Ebro-1944) falls at one end of the latter cluster, on an axis of variation shared by strains from Mexico, Brazil, Colombia and Peru.

**Fig. 1:**
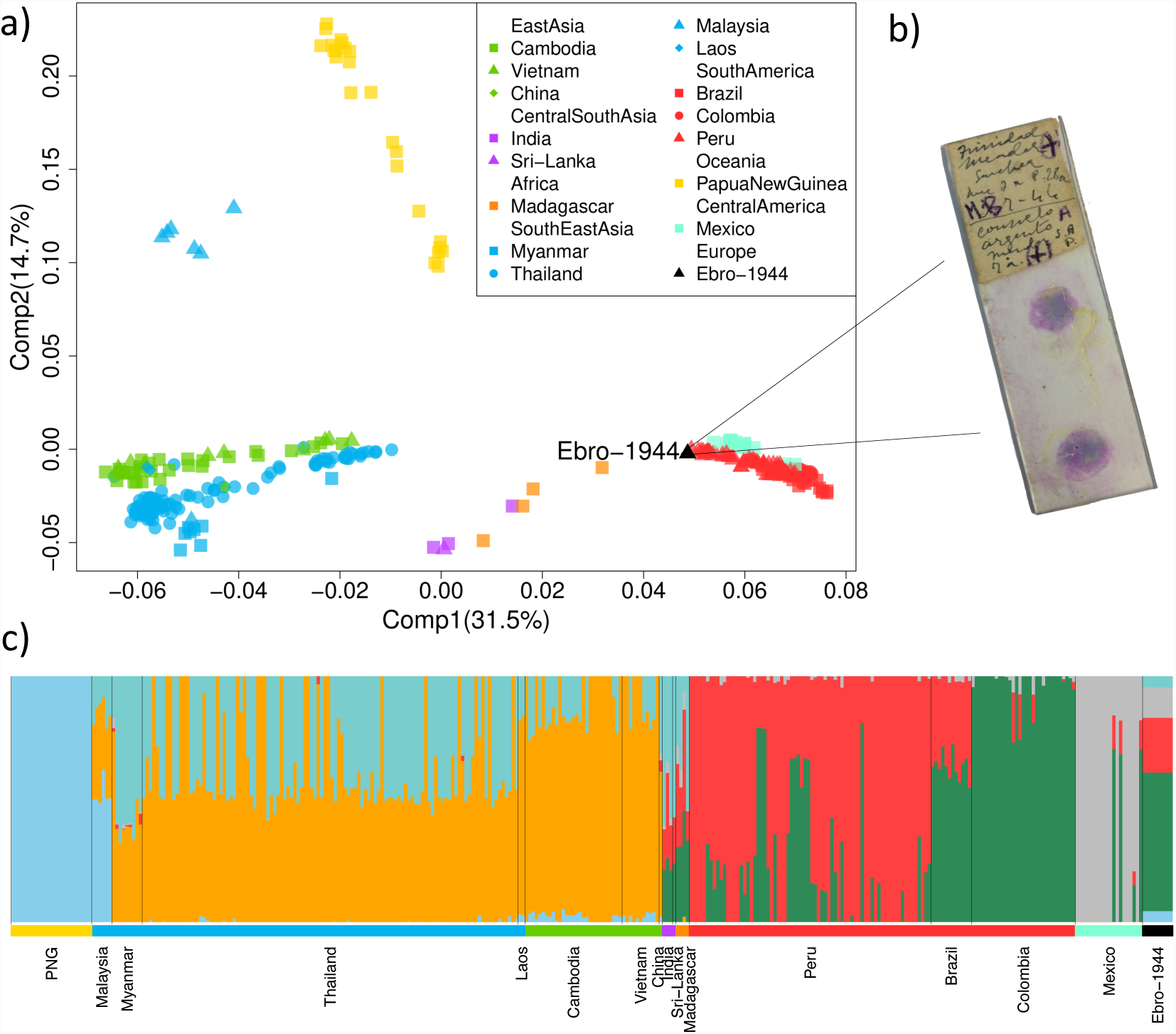
a) Principal components analysis (PCA) of the historic Ebro-1944 sample together with a geographically diverse set of modern *P. vivax* strains. b) Example microscopy slide stained with the blood of patient’s infected with malaria from the Ebro Delta, Spain, in the 1940s. c) Unsupervised ADMIXTURE clustering analysis at *K*=6. Samples are arranged by geographic region and coloured as in a).

Model-based clustering implemented in an unsupervised ADMIXTURE analysis (fig. 1c, Supplementary fig. 6) provided qualitatively consistent inferences to those observed by PCA, with one ancestry component maximised in Oceanian samples, two largely shared by East and South East Asian samples and three further components which largely differentiate samples from South and Central America. The ancestry of Ebro-1944 is mostly modelled by the three components identified in samples from South America (~56% Peru-like and ~22% Colombia-like) and Central America (12% Mexico-like), although a minor proportion of ancestry is shared with both South East Asian (~5%) and Oceanian (~5%) samples.

To formally test these suggested relationships, we calculated *f*4 statistics (Patterson et al. 2012) of the form (*P. cynomolgi*, Ebro-1944; X, Y), where X and Y are tested for all combinations of samples isolated from 15 worldwide locations (fig. 2). *P. cynomolgi* was selected as an outgroup as it represents the closest non-human infecting *Plasmodium* species (Tachibana et al. 2012). This statistic is designed to quantify the covariance in allele frequency differences between *P. cynomolgi* and Ebro-1944 relative to *P. vivax* from sampled worldwide locations (X and Y), with a more positive *f*4 value indicating a closer relationship of Ebro-1944 to samples from Y relative to X. Using this framework and testing all possible topological relationships, Ebro-1944 was found to share significantly more derived alleles with Central and Southern American strains compared to those sampled from South East and East Asia (fig. 2, Supplementary fig. 7). These results collectively support the presence of a cline of ancestry stretching from Europe to the Americas, with our eradicated European strain showing strong phylogenetic affinity to modern *P. vivax* samples from Mexico, Brazil and Peru.

**Fig. 2:**
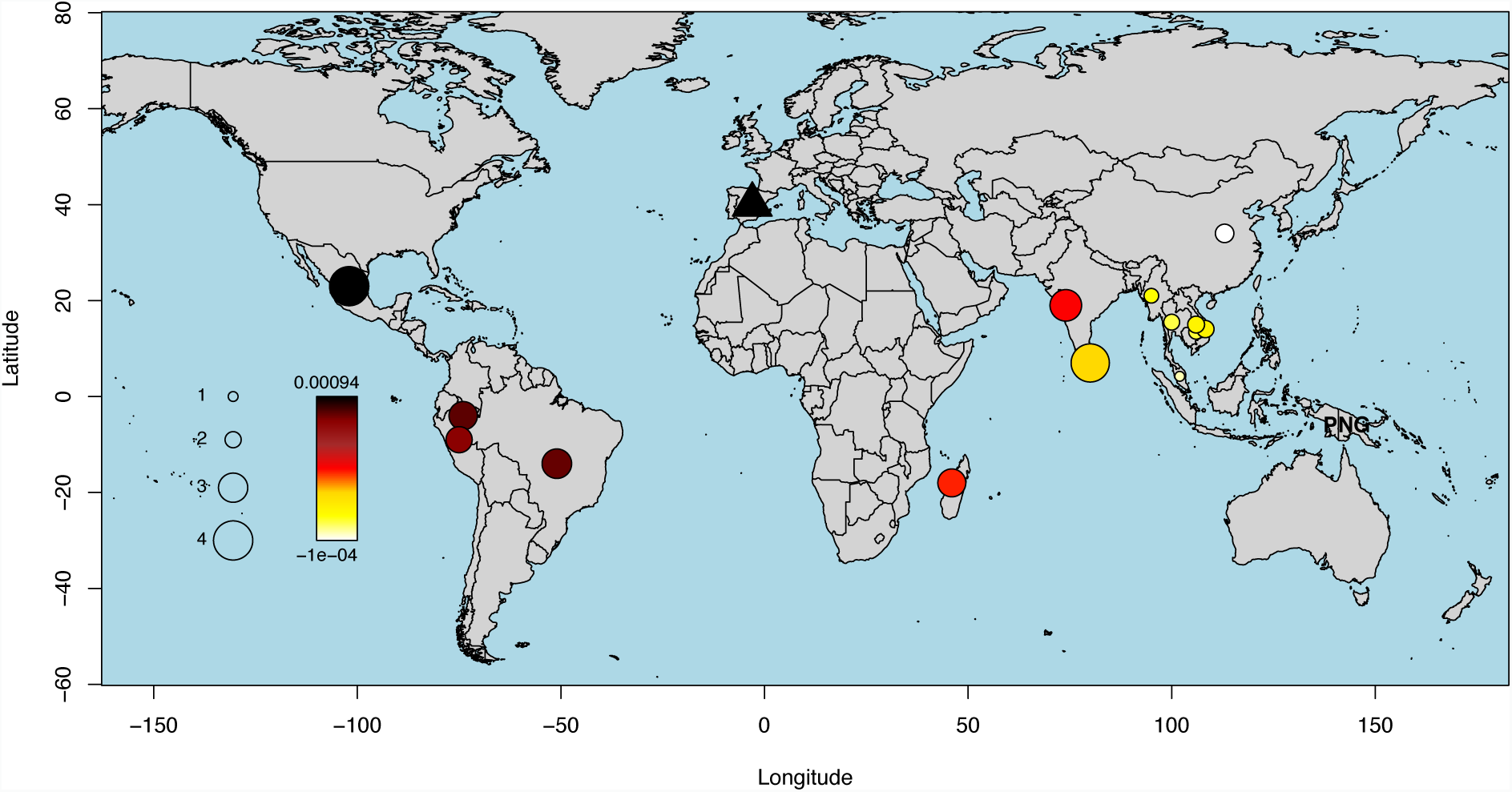
*f*4-values inferred under the test relationship (*P. cynomolgi*, Ebro-1944; Papua New Guinea (PNG), Y), where Y iterates through the geographic sampling locations of our included strains. The colour scale provides the value of the *f*4 statistic with the significance (absolute z score), assessed through block jack-knife resampling, provided by the circle size. A more positive *f*4 value indicates a closer relationship of Ebro-1944 to Y relative to PNG.

Additional evidence was also obtained using an unrelated method designed to cluster global samples based on inferred patterns of haplotype sharing (Lawson et al. 2012). Considering haplotype variation rather than allele frequency differences increases power to resolve fine-scale genetic structure (Leslie et al. 2015) and is robust to potential SNP calling errors (Conrad et al. 2006). Haplotype-based clustering grouped Ebro-1944 with samples from South and Central America (fig. 3a). This result was robust to the inclusion of different samples in the dataset, to moderate imputation, and remained consistent when either uncorrelated sites or linked sites were considered (Supplementary fig. 8-10, Supplementary Section 4 and Methods).

**Fig. 3:**
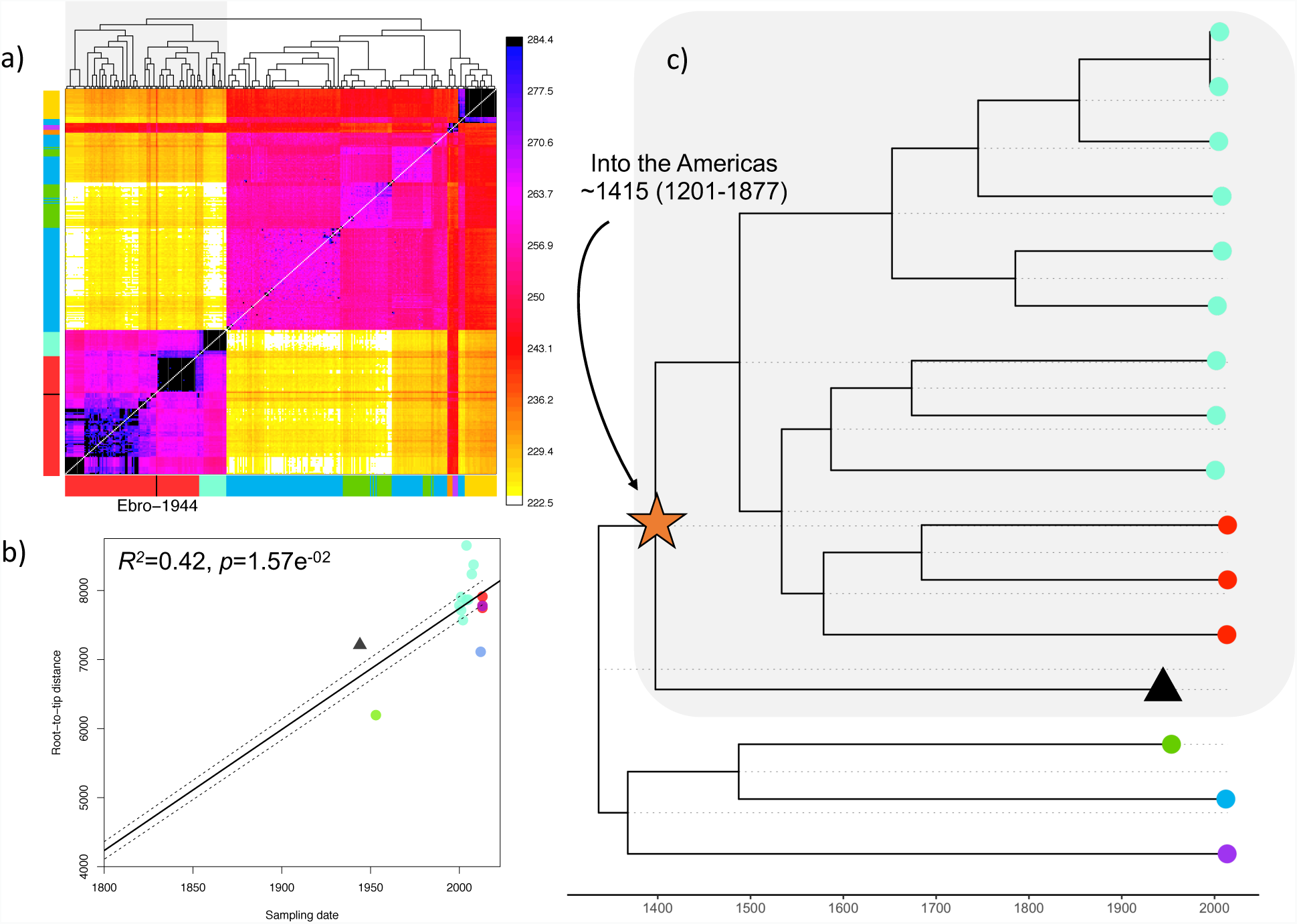
a) CHROMOPAINTER’s inferred counts of matching DNA genome wide that each of the 104 inferred clusters (columns) is painted by each of the 104 clusters (rows). The tree at top shows fineSTRUCTURE’s inferred hierarchical merging of these 104 clusters and the colours on the axes give the continental region and population to which strains in each cluster are assigned. Ebro-1944 is depicted in black and clusters with the sample from Peru and Brazil. b) Root-to-tip distances of our included *P. vivax* strains correlated with the date of isolation. The regression was significant following 1000 random permutations of sampling date. c) Tip-dated phylogenetic tree obtained with BEAST 2. The mean posterior probability for time to the most recent common ancestor of the split between the historical European strain and the American isolates is indicated. Strains are coloured as in fig. 1a.

Given the strong phylogeographic affinity of Ebro-1944 to *P. vivax* currently in circulation in the Americas, we tested whether the divergence of Ebro-1944 from American strains is better explained by a deep-split, for example at the time of the original human settlement of the Americas at least 15,000 years ago, or is more consistent with a recent introduction to the Americas. Genomes obtained from historical or ancient materials provide unique opportunities to calibrate phylogenetic trees, by directly associating sampling dates with the sequences representing the phylogeny tips (terminal nodes). These in turn enable inference of divergence times and mutation rates without the need for any other age-related external data (Rieux and Balloux 2016). Therefore, to infer the temporal relationship of Ebro-1944 to strains from the Americas, we included our historic sample together with 15 closely-related publicly available genomes sampled over a range of time periods (Supplementary Table 4). We selected strains predominantly from the Americas, with three additional genomes included from India, Myanmar and North Korea to root the topology and increase the time span of our dataset.

To account for the possible confounding effect of genetic recombination, we filtered the resulting alignment for a set of high confidence congruent SNPs. Specifically, we removed all homoplasies i.e. SNPs in the alignment in conflict with the maximum parsimony phylogeny (see Methods). This approach identifies, with no required prior knowledge, regions of the genome that are hyper-variable, deriving from recombination or mixed infections, as well as filtering SNPs which may have been erroneously called due to low sequence quality or post-mortem damage. The resulting alignment exhibited a significant positive correlation between the root-to-tip phylogenetic distances of a maximum likelihood phylogeny and the time of sampling, indicating the presence of detectable temporal accumulation of *de novo* mutations within the timescale of our dataset (fig. 3b).

Mutation rates were subsequently estimated using the Bayesian phylogenetic tool BEAST2 (Bouckaert et al. 2014) testing a range of demographic and clock rate priors. We estimated the mutation rate over the tested alignment to 5.57E^−7^ substitution/site/year [HPD 95% 2.75E^−8^ – 1.06E^−6^]. Though a broad estimate, we obtained low values (<0.15) of the standard deviation of the uncorrelated log-normal relaxed clock (ucld.stdev), suggesting little variation in rates between branches. This approximation enabled us to infer that the historical Ebro-1944 genome shares a common ancestor with strains in the South-American cluster dating to the 13^th^ to 19^th^ centuries (mean 1415; HPD 95% 1201-1877CE) (fig. 3c, Supplementary fig. S11, Supplementary Table 5).

A number of mutations conferring resistance to antimalarial drug treatments developed in the later decades of the 20th century have been identified in *P. vivax*. For instance, mutations in the *pvdhfr* gene are known to be involved in resistance to pyrimethamine (de Pecoulas et al. 1998; Imwong et al. 2001; Imwong et al. 2003; Huang et al. 2014) whilst mutations at the *pvdhps* gene confer resistance to sulfadoxine (Korsinczky et al. 2004; Menegon et al. 2006; Hawkins et al. 2009). Other genes, including *pvmdr1* (Brega et al. 2005; Sá et al. 2005; Barnadas, Tichit, et al. 2008), *CRT* (Suwanarusk et al. 2007) or *pvmrp1* (Dharia et al. 2010) are thought to be involved in chloroquine resistance. *P. vivax* populations also exhibit high genetic diversity in genes related to immune evasion and host infectivity, including *MSP10*, *MSP7* and *CLAG*. We annotated 4,800 SNPs in the Ebro-1944 genome (Supplementary Table 6). Of these, 1195 are missense mutations, which represent 60.6% of the genic mutations. Similar ratios have been reported in other *P. vivax* strains (Hupalo et al. 2016; Pearson et al. 2016; de Oliveira et al. 2017).

Ebro-1944 carries the derived allele in three SNPs located in genes functionally associated to antimalarial drug resistance. Two of these, Val1478Ile and Thr259Arg (Dharia et al. 2010; de Oliveira et al. 2017) are in the *pvmdr1* gene and another, Met205Ile (Hawkins et al. 2009), occurs in the *pvdhps* gene (Supplementary Tables 7-8). We also identified a previously undescribed mutation (Leu623Arg) in *CLAG*, a gene associated with host infectivity (Gupta et al. 2015). Additionally, we screened multiple loci (N=516) that have previously been reported to exhibit strong signals of recent natural selection in a geographically diverse set of modern *P. vivax* samples (Hupalo et al. 2016). Some of these regions encompass genes previously known to be involved in anti-malarial drug resistance, including three with strong experimental validation of resistance phenotypes: *pvmdr1*, *dhfr* and *dhfps* (Haldar et al. 2018). Our historical genome has 355 of these positions covered by at least two reads, of which Ebro-1944 exhibits the ancestral allele in 349 (Supplementary Table 9). Therefore, only six of the 355 derived variants with high *F*_ST_ values (including the previously mentioned Met205Ile variant at *pvdhps* gene) were present in the historical European sample, suggesting a rapid accumulation of resistance conferring mutations in more recent strains.

## Discussion

Our genome-wide analyses of a historic European *P. vivax* nuclear genome confirm the inference, previously based solely on mtDNA, that extinct European *P. vivax* are closest genetically to strains currently in circulation in Central and South America (Gelabert et al. 2016). Historical accounts of the presence of tertian (*P. vivax*) malaria in Europe date to at least Classical Greece (De Zulueta 1973; Carter 2003), suggesting that the most parsimonious explanation is an introduction of *P. vivax* from Europe into the Americas and not the other way around. A migration event from Europe into the Americas is further supported by the European Ebro-1944 *P. vivax* strain falling as an outgroup to all the strains from the Americas (fig. 3c). Finally, the estimated mean age of divergence in the 15^th^ century between Ebro-1944 and strains in circulation in the Americas (fig. 3c) is consistent with an introduction of *P. vivax* malaria into the Americas by European colonists.

Whether any agent of malaria was present in the Americas prior to European contact (1492) has been debated (Carter 2003; Hume 2003; de Castro and Singer 2005). It is relatively unlikely that any malarial parasites would have survived the journey across the Bering strait during the initial peopling of the Americas some 15,000 years ago (Waters et al. 2018) due to the absence of mosquito vectors at these high latitudes allowing the parasites to fulfill their life cycles (Tanabe et al. 2010). However, accounts of the therapeutic use by Incas of cinchona tree bark, from which quinine derives, have been interpreted as suggestive of malaria being present in the Americas pre-Columbian contact (Escardo 1992). Though, its use may have been motivated by the effectiveness of quinine in the treatment of other fever-causing illnesses.

Older split times between *P. vivax* lineages from the Americas and the rest of the world than the one we inferred based on whole genome sequences have been suggested from the analysis of mtDNA genomes (Taylor et al. 2013; Rodrigues et al. 2018). Though, we note that, even when accounting for the posterior uncertainty around our inferred mutation rate, all possible temporal estimates place the split time between the European and American strains as incompatible with an introduction of *P. vivax* into the Americas alongside the first humans to colonize the continent. Our results are therefore highly supportive of an introduction of *P. vivax* to the Americas during the European colonial period, with our range also consistent with the transatlantic slave trade between Africa and Spanish and Portuguese-run ports in Central and South America. Our analyses of the nuclear genome further point to a minor genetic component in American strains shared with strains from Madagascar, India and Sri Lanka (fig. 1c, fig.3a). We interpret this as likely evidence for secondary genetic introgression into the American *P. vivax* population by lineages from different regions of the world, which would be consistent with the high *P. vivax* mtDNA diversity previously described in the Americas (Taylor et al. 2013; Hupalo et al. 2016; de Oliveira et al. 2017; Rodrigues et al. 2018).

In this work, we restricted the phylogenetic dating to the migration of *P. vivax* into the Americas. There is currently no consensus over the age of the most recent common ancestor of all extant, worldwide *P. vivax* strains. Some previous estimates inferred the origin of *P. vivax* to around 5,000 or 10,000 years ago (Carter 2003; Leclerc et al. 2004; Lim et al. 2005), whereas far older dates have been proposed, including 45,000-81,000 years (Escalante et al. 2005) or even 53,000-265,000 years (Mu et al. 2005). An extrapolation of our inferred evolutionary rates to the global diversity of extant *P. vivax* strains would point to a recent origin for the parasite. Though, formally estimating the age of *P. vivax* through a phylogenetic ‘tip dating’ approach poses a series of analytical challenges. For example, the homoplasy screening method we employed to exclude homoplasies caused by mixed infections and likely genetic recombination is currently not computationally tractable for a dataset comprising a large number of globally sourced whole *P. vivax* genomes. Possible solutions to this problem may arise through optimisation of sequencing protocols to generate higher quality Plasmodium whole genome sequences together with the development of downstream bioinformatics approaches designed to propagate genotype calling uncertainty and deconvolve mixed infections (Zhu et al. 2018). This should be aided by the generation of further high-quality reference genomes across the Plasmodium genus, for example using long-read sequencing technology (Auburn et al. 2016; Pasini et al. 2017; Gilabert et al. 2018; Otto et al. 2018).

Reconstruction of the phylogeographic relationships between *P. vivax* strains is complicated further by mounting evidence for the zoonotic potential of *P. vivax* and *P. vivax-like* strains; with host-jumps likely having occurred several times in the parasite’s evolutionary history (Prugnolle et al. 2013; Liu et al. 2014; Liu et al. 2017; Loy et al. 2017). The discovery of the platyrrhine protozoa, *Plasmodium simium*, as morphologically (Ott 1967) and genetically (Leclerc et al. 2004; Escalante et al. 2005; Lim et al. 2005) indistinguishable from *P. vivax* suggests very recent host transfers between South American monkeys and humans; in some cases responsible for the incidence of zoonotic malarial disease (Brasil et al. 2017; Buery et al. 2017). This raises important questions as to what should be considered as the host-range of *Plasmodium vivax* and queries what species delineation should be applied in comparative genomics studies. Recurrent host jumps of *P. vivax* lineages into humans from animal reservoirs, with subsequent demographic expansions and possible lineage replacements, may point to a recent age for the ancestor of all extant *P. vivax* strains, despite now extinct lineages of the parasite having likely plagued humans for far longer.

Malaria is widely believed to have exerted one of the strongest selective forces on the human genome (Hedrick 2012). Well known examples of selection against malaria include protective mutations at the *HBB* gene that give rise to resistant isoforms of proteins such as HbS and HbE in African and Asian populations, respectively, and mutations at the *G6PD* gene which are broadly spread in African populations and are also present in the Mediterranean (Tishkoff et al. 2001; Kwiatkowski 2005; Howes et al. 2012). One of the best-known examples of directional selection is the FY*0 Duffy blood negative genotype that confers resistance to *P. vivax* which is close to fixation in sub-Saharan Africa but essentially absent in other regions of the world. The protein is the key invasion receptor for the human malarial parasites *P. vivax*, *P. knowlesi* and the simian malarial parasite *P. cynomolgi* (Kosaisavee et al. 2017). Exposure to an ancestor of *P. vivax* may have led to the sweep of the FY*0 allele in sub-Saharan Africa some 40,000 years ago and may represent the fastest known selective sweep for any human gene (McManus et al. 2017).

Though, given the relatively mild clinical symptoms of modern *P. vivax*, it is not inconceivable that the selective forces that led to the fixation of the FY*0 allele in sub-Saharan Africa may have been caused by another malaria parasite. A further complicating factor stems from increasing evidence accumulating for previously unrecognised endemic *P. vivax* circulating in human populations from Sub-Saharan Africa, including in Duffy-negative hosts (Gunalan et al. 2018; Twohig et al. 2019). The presence of endemic *P. vivax* malaria in sub-Saharan Africa may suggest that the FY*0 allele may confer only partial protection against *P. vivax* malaria. Our results, which point to a recent, post-European contact exposure of Native American populations to *P. vivax* malaria, would not have had the time to drive the emergence and spread of resistance alleles comparable to those observed in Africa and Europe. Consistently, to date, no known malaria resistance variants have been identified in Native Americans (Hume 2003; Kwiatkowski 2005).

Irrespective of its role in the past, the rapid spread of *P. vivax* strains resistant to antimalarial drugs is an area of increasing concern. Initially, Chloroquine was established as the main therapy against *P. vivax* infections in 1946 (Most and London 1946; Baird 2004). It was a well-tolerated and effective treatment until resistance appeared in the late 1980s and spread through the entirety of the endemic range of *P. vivax* (Rieckmann et al. 1989). However, despite extensive drug resistance within present-day *P. vivax* populations, caused by a variety of resistance loci in multiple genes (Hupalo et al. 2016), chloroquine-primaquine combined therapy remains the most commonly prescribed treatment (Phillips et al. 1996), as few other therapeutic strategies are available. The increase in frequency of drug resistant strains is thus a significant public health threat with major human and economic costs.

Our historical sample predates the use of all anti-malaria drugs, with the exception of quinine which was introduced in Europe as early as 1683 (Achan et al. 2011). Ebro-1944 carries the ancestral allele in an overwhelming number of SNPs (99.3%) known to have undergone selection in modern strains, including those associated with drug resistance in genes such as *DHFR-TS* (de Pecoulas et al. 1998; Leartsakulpanich et al. 2002; Imwong et al. 2003; Ganguly et al. 2014; Huang et al. 2014) and *MDR1* (Brega et al. 2005; Sá et al. 2005; Barnadas, Ratsimbasoa, et al. 2008; Orjuela-Sánchez et al. 2009) (Supplementary Table 8). Conversely, Ebro-1944 carries three variants in the *pvmdr1* and *pvdhps* genes that are plausible drug-resistance candidates against sulfadoxine and chloroquine. The presence of these alleles in Ebro-1944 might reflect standing variation in historical *P. vivax* populations for alleles providing resistance to modern antimalarial drugs. Alternatively, these could have been selected for by the historical use of quinine.

Our study stresses the value of old microscopy slides and, more generally, of antique medical collections, as a unique and under-used resource for retrieving genomic information on pathogens from the past, including eradicated strains that could not be studied from contemporary specimens. For example, in addition to the results reported here, we also retrieved a partial *P. falciparum* genome from the same set of slides, which allowed us to demonstrate a stronger phylogeographic affinity of the extinct European *P. falciparum* lineage to present-day strains in circulation in central south Asia, rather than Africa (de-Dios et al. 2019 *in press*). We note that the slides we analysed here were stained but not fixed and it remains to be explored what additional DNA damage is exerted by different fixation methods. Additionally, our slides date from the years 1942-1944; however, it is likely that older slides are available in both public and private collections given the popularity of microscopy in Victorian times. A future objective will be to ascertain if massive genomic data retrieval can be achieved from even older microscopy slides.

There is also potential to retrieve ancient *Plasmodium* sequences directly from archaeological specimens. The recent retrieval of *P. falciparum* sequences from ancient Roman human skeletal remains (Marciniak et al. 2016) demonstrates this approach is technically feasible, and as *P. vivax* infection is more prevalent than *P. falciparum*, it is plausible that further ancient strains from osteological material could be reported in the near future. An additional possibility would be to directly retrieve *Plasmodium* sequences from *Anopheles* remains preserved in ancient lake sediments or in museum collections. The generation of additional historical sequences - both from Europe and from the Americas - together with an increased sequencing effort of extant *P. vivax* strains from under-sampled areas is our best hope to reconstruct the evolutionary history of this major parasite in more detail.

## Materials and Methods

### Samples

The slides analysed here belong to the personal collection of the descendants of Dr. Ildefonso Canicio, who worked in the antimalarial centre established by the Catalan Government at Sant Jaume d’Enveja (Ebro Delta, Spain) in 1925. Four slides were analysed, three of them included in a previous study (Gelabert et al. 2016). The new sample was a drop of blood from a double slide, stained with Giemsa (fig. 1b).

### DNA extraction

DNA extraction was performed by incubating the slide with 20μL of extraction buffer (10 mM Tris-HCl (pH 8), 10 mM NaCl, 5 mM CaCL, 2.5 mM EDTA, 1% SDS, 1% Proteinase K, 0.1% DTT (w/v)) in an oven at 37°C for 20 minutes for a total of three rounds. The resulting dissolved bloodstain and buffer were collected in a 1.5 mL Lobind Eppendorf and then incubated for one hour at 56ºC. This was subsequently added to 10x volume of modified binding buffer (Allentoft et al. 2015) and passed through a Monarch silica spin column (NEB) by centrifugation (Supplementary Methods Section 1 and Supplementary fig. 12). The column was washed once with 80% ethanol and DNA was subsequently released with EBT buffer to a final volume of 40μL (see Supplementary Material Section 1). All the analyses were performed in dedicated ancient DNA laboratories where no previous genetic work on *Plasmodium* had been carried out, both in Barcelona (extraction of slides in 2016) and Copenhagen (extraction of slides in 2017).

### Library preparation and DNA sequencing

Shotgun sequencing libraries for the Illumina platform were prepared using a single-tube protocol for double-stranded DNA (Christian et al. 2017), with minor modifications and improvements as detailed in Mak *et al.* (2017) (Supplementary Methods Section 2). Sequencing was performed at the Natural History Museum of Denmark on one lane of an Illumina Hiseq 2500 instrument in paired end mode running 125 cycles.

### Sequence mapping

The sequenced reads were analysed with FastQC to determine the quality prior to and after adapter clipping. The 3’ read adapters and consecutive bases with low quality scores were removed using cutadapt 1.18 (Martin 2011). Reads shorter than 30 bp and bases with a quality score lower than 30 were also excluded. To increase the final coverage, all *Plasmodium* reads were pooled and an in-house script was used to discriminate reads mapping more confidently to the *P. vivax* (PvP01) (Auburn et al. 2016) compared to the *P. falciparum* 3D7 (Gardner et al. 2002) reference genomes based on edit distance. Mapping of *P. vivax* reads was then performed with Burrows-Wheeler Aligner (BWA) (Li and Durbin 2009) 0.7.1 aln (Supplementary Methods Section 2). Duplicated reads were deleted using Picard tools 2.18.6 MarkDuplicates. MapDamage 2.0 (Ginolhac et al. 2011) was applied to check for signatures of post-mortem damage at the ends of the reads to validate the reads were associated with a historic sample rather than deriving from modern contamination. C to T and G to A substitutions at the 5’ ends and 3’ ends, respectively, were found to be present at a frequency of about 2.5% (Supplementary fig. 1); consistent with the age of the sample and in agreement with the degree of damage previously detected in the mtDNA reads (Gelabert et al. 2016). As a result, the first two nucleotides of each read were also trimmed.

The newly generated paired-end *P. vivax* reads were merged with the sequencing reads generated in 2016 (Gelabert et al. 2016). We refer to the resulting pooled sample as Ebro-1944. Genotypes were called from the alignment with GATK v3.7 UnifiedGenotyper (McKenna et al. 2010), using a minimum base quality of 30 and the standard confidence call threshold of 50. Genotype calls were filtered further with VCFtools (Danecek et al. 2011), excluding: i) heterozygous calls, ii) calls with depth of coverage <2, iii) calls present in the telomeres and subtelomeres (Pearson et al. 2016), iv) indels (Supplementary fig. 2 and 3). SNPs were annotated using SnpEff (Cingolani et al. 2012) for the analysis of variants associated with drug resistance.

### Population genetics dataset

The dataset used in population genetics analyses comprised the nuclear sequences of 337 previous published samples of *P. vivax* (Hupalo et al. 2016; Pearson et al. 2016; Cowell et al. 2017; Cowell et al. 2018; Rodrigues et al. 2018), representing the global diversity of currently available *P. vivax* genomes (Supplementary Tables 3 and 10).

Sequence reads were aligned against the Sal1 reference genome (Carlton et al. 2008) with BWA 0.7.1 aln (Li and Durbin 2009) using default parameters. Duplicated reads were removed using Picard tools 2.18.6. Reads with mapping qualities below 30 were removed using SAMtools 1.6 (Li et al. 2009). For analyses requiring incorporation of an outgroup, we additionally mapped reads from the *P. cynomolgi* M strain (Pasini et al. 2017) against the Sal1 reference genome following the same protocol.

We used GATK v3.7 UnifiedGenotyper (McKenna et al. 2010) for SNP calling with a number of adjustments for working with *Plasmodium* genomes and ancient DNA. First, we selected 297 samples that had more than the 70% of the Sal1 reference genome covered and presented more than 3000 substitutions. Using this filtered dataset of high-quality samples, we called variants using a mapping quality >30, depth of coverage >20 and a standard call confidence >50. We removed those SNPs that mapped to repetitive regions of the *P. vivax* reference genome (Pearson et al. 2016), heterozygous calls suggesting possible mixed infections, and SNPs that were present in less than three samples. The resultant dataset included 131,309 SNPs and 277 strains. We used this catalogue to genotype all remaining samples in the dataset. For the individual genotyping, we called SNPs with GATK v3.7 UnifiedGenotyper as before, removing all heterozygous calls, and all variants with a minor allele frequency (MAF) below 0.01%. The final dataset comprised 338 samples and 128,081 SNPs. The genotypes for Ebro-1944 were called by selecting one random read that mapped each position of the dataset (Mathieson et al. 2015). This resulted in the Ebro-1944 sequence covering 77,425 of the 128,081 total included positions.

### Allele-frequency based measures of population structure

The population genetics dataset was filtered for SNPs in high linkage disequilibrium (LD) using a 60 SNP sliding window, advancing each time by 10 steps, and removing any SNPs with a correlation coefficient ≥0.1 with any other SNP within the window (Chang et al. 2015). This left a pruned dataset of 38,358 SNPs for analyses relying on independent SNPs. We applied PCA to the LD pruned global dataset restricted to only sites covered in all samples (Chang et al. 2015). We additionally clustered our dataset using the unsupervised clustering algorithm ADMIXTURE 1.3.0 (Alexander and Novembre 2009) for values of K between 1-15. K=6 provided the lowest cross-validation error (Supplementary fig. 6, fig. 1c). To evaluate the relationship of Ebro-1944 to other global strains we calculated *f*4 statistics using qpDstat available within AdmixTools (Patterson et al. 2012). Setting Ebro-1944 as a target, we explored which strains, grouped by geographic label, share more alleles with Ebro-1944 relative to every pairwise combination of modern strain(s) (X and Y) in our reference dataset and relative to *P*. *cynomolgi* as an outgroup: *f*4(*P. cynomolgi*, Ebro-1944; X, Y).

### Inferring patterns of allele and haplotype sharing

In addition, we applied an unrelated method to explore patterns of allele and haplotype sharing implemented in CHROMOPAINTER v2 (Lawson et al. 2012). Unlike *f*-statistics, this approach does not rely on a user specified topology and can thus consider the relationship of all strains to all others collectively. As this approach requires low levels of missingness across comparisons, we filtered the previously described unpruned population genetics dataset for only the positions present in Ebro-1944 and retained only those samples with ≤10% missing data (77,420 sites, 218 samples). A schematic of the workflow is provided in Supplementary fig. 5.

Briefly, CHROMOPAINTER calculates, separately for each position, the probability that a “recipient” chromosome is most closely related to a particular “donor” in the dataset under a copying model framework (Li and Stephens 2003). Here, we use all strains as donors and the equivalent strains as recipients in an “all-versus-all” painting approach. To cluster strains, fineSTRUCTURE (Lawson et al. 2012) was applied to the all-versus-all coancestry matrix to group strains based on their inferred painting profiles. Given the variable missingness across the modern strains included in the alignment we implemented several analyses using the CHROMOPAINTER unlinked (allele-sharing) implementation, as well as under the linked CHROMOPAINTER (haplotype-sharing) model. To use the latter we performed various levels of imputation followed the protocol set out by Samad et al (2015) (Samad et al. 2015) for *P. falciparum* in BEAGLE v3.3.2 (Browning and Browning 2013). The consistency of our inference under different imputation and filtering criteria was assessed by linear regression (Supplementary fig. 10). Further details are provided in Supplementary Section 4.

### Drug resistance variants analysis

We used the annotations provided by SnpEff to identify non synonymous mutations in genes previously described as being related to antimalarial drug resistance and host infectivity (Cingolani et al. 2012; Gupta et al. 2015; Hupalo et al. 2016; Pearson et al. 2016; Rodrigues et al. 2018). We also screened a set of previous described positions that have shown recent signals of selection in *P. vivax* populations (Supplementary Table 7). In addition, we screened all potential genetic variants found in Ebro-1944, comprising more than 4000 SNPs (Supplementary Table 9).

### Phylogenetic analysis and dataset

The whole-genome sequences of 15 modern *P. vivax* samples and Ebro-1944 were mapped against the PvP01 reference assembly (Auburn et al. 2016) (Supplementary Table 4). These strains were selected with a focus on the Americas but also to include strains sampled over a large temporal span as is required for phylogenetic tip-dating. The main motivation for using the PvP01 reference genome for mapping sequence data for phylogenetic analyses stems from this assembly offering a better definition of sub-telomeric genes and repetitive *pir* genes that usually lie in recombinant regions, thus providing greater power to identify and exclude these parts of the genome (Auburn et al. 2016). After calling variants with GATK version 3.7 UnifiedGenotyper, polymorphisms were further filtered by selecting only those SNPs with a coverage >1 an average mapping quality >30, a genotype quality >30 and by removing heterozygous positions. The exonic positions were then classified as synonymous and non-synonymous with SnpEff (Cingolani et al. 2012).

Our final phylogenetic timeline dataset spanned 69 years (1944-2013) of evolution (Supplementary Table 4) across a 34,452 SNP alignment. In order to filter this alignment for only congruent SNPs for phylogenetic dating we first generated a maximum parsimony phylogeny in MEGA7 (Kumar et al. 2016), evaluating support for each branch over 100 boot-strap iterations. We then screened for homoplasies, sites in the 34,452 SNP alignment that do not support the maximum parsimony phylogeny, using HomoplasyFinder (Crispell et al. 2019). This led to the identification of 13,112 homoplastic SNPs, many of which fell in sub-telomeric regions. All homoplastic sites were subsequently removed from the alignment. In this way we screen and exclude variants from the original alignment that fall in hyper-variable regions, that may arise from inaccurate SNP calling or post-mortem damage, as well as removing regions that derive from between-lineage genetic recombination or mixed infections.

### Estimating a timed phylogeny

To investigate the extent of temporal signal existing in our homoplasy cleaned timeline alignment, we built a maximum-likelihood phylogenetic tree, without constraining tip-heights to their sampling times, using RaxML (Stamatakis 2014). After rooting the tree on the *P. cynomolgi* genome, we computed a linear regression between root-to-tip distance and sampling time using the roototip function from BactDating (Didelot et al. 2018). To further confirm the presence of a significant temporal signal, we assessed the significance of this regression following 1000 steps of date randomisation (fig. 3b). After confirmation of temporal signal in the dataset, substitution rates were estimated by running a tip-calibrated inference using Markov chain Monte Carlo (MCMC) sampling in BEAST 2 (Bouckaert et al. 2014). The best-fit nucleotide substitution model was estimated as TN93 following evaluation of all possible substitution models in BModelTest (Bouckaert and Drummond 2017). To minimise prior assumptions about demographic history we tested three possible demographic models: the coalescent constant, coalescent exponential and coalescent Bayesian skyline. In each case we set a log normal prior on a relaxed evolutionary clock as well as testing under a strict clock model.

To calibrate the tree using tip-dates only, we applied flat priors (i.e., uniform distributions) for the substitution rate (1.10E12 - 1.10E2 substitutions/site/year), as well as for the age of any internal node in the tree. We ran five independent chains in which samples were drawn every 50,000 MCMC steps from a total of 500,000,000 steps, after a discarded burn-in of 10,000,000 steps. Convergence to the stationary distribution and sufficient sampling and mixing were checked by inspection of posterior samples (effective sample size >200) in Tracer v1.6. The best-fit model was selected based on evaluation of both the median likelihood value of the model and the maximum likelihood estimator following path sampling (Baele et al. 2012) (Supplementary Table S5).

## Acknowledgements and funding information

We are grateful to Thomas D. Otto (University of Glasgow) and Thomas Lavsten (University of Copenhagen) for helpful comments and suggestions and to the descendants of Dr Canicio, Miquel and Ildefons Oliveras for sharing with us their slides. We additionally would like to thank Dr. George Busby and Dr. Jacob Almagro for useful discussions on chromosome painting of Plasmodium *sp*. This research was supported by a grant from Obra Social “La Caixa”, Secretaria d’Universitats i Recerca Programme del Departament d’Economia i Coneixement de la Generalitat de Catalunya (GRC 2017 SGR 880) and by FEDER-MINECO (PGC2018-095931-B-100) to C.L.-F and an ERC Consolidator Grant (681396-Extinction Genomics) to MTPG. LvD and FB acknowledge financial support from the Newton Trust UK-China NSFC initiative (grants MR/P007597/1 and 8166113800). *P. vivax* genomes are deposited at the European Nucleotide Archive under accession numbers XXXX-XXXX. We also thank the Danish National High Throughput Sequencing Centre for help in sequencing.

## Author contributions

P.G., M.T.P.G., L.v.D., F.B. and C.L.-F. conceived and designed the study; R.E. and C.A. discovered the slides; C.C. developed and performed laboratory analysis; L.v.D., P.G., A.R., T. d.-D., S.G., R.F., I.O., M.d.M. analysed data and performed computational analyses; S.H., F.B., T.M.-B. and I.M. provided comments and suggested analyses; L.v.D., F.B., and C.L.-F. wrote the paper with inputs from all co-authors.

