## Supplementary Information for "*Plasmodium vivax* Malaria viewed through the lens of an eradicated European strain"

### CONTENTS

1. DNA extraction from glass slides
2. DNA Sequencing and mapping
    2.1 Shotgun library preparation for Illumina
    2.2 Quantitative PCR (qPCR)
    2.3 Index PCR amplification
    2.4 Genome mapping
3. Human reads and contamination
4. Allele and haplotype sharing analyses

SUPPLEMENTARY FIGURES

SUPPLEMENTARY TABLES

REFERENCES

### 1. DNA extraction from glass slides

To improve retrieval of DNA from glass slides, a modified protocol relative to Gelabert et al. 2016 was developed. A number of extraction rounds were performed by applying 20 µL of extraction buffer (10 mM Tris-HCl (pH 8), 10 mM NaCl, 5 mM CaCL, 2.5 mM EDTA, 1 % SDS, 1% Proteinase K, 0.1% DTT (w/v)) to the bloodstain on the slide. The slide was then immobilized horizontally in a 50 mL Falcon tube with tin foil and incubated in an oven at 37°C for 20 minutes. This process was repeated three times, and each time the incubated extraction buffer was collected in the same 1.5 mL Lobind Eppendorf tube using a 20 µL pipette. The collected buffer was then incubated for one hour at 56°C on a heatblock. Subsequently 10x volume of modified binding buffer was added (Allentoft et al. 2015) and passed through a Monarch silica spin column (NEB) at 3000xg.

Using a separate approach, we tested Monarch columns, which allow retrieval of slightly more (~8%) short DNA compared to Qiagen Minelute spin columns (see **Supplementary Figure 12**). The column was then added to 800 µL 80% ethanol, spun at 10.000xg for one minute, followed by three minutes at 17.000xg to spin dry. 20 µL of EBT buffer was added to the column and incubated for 10 minutes at 37°C on a heatblock before being released into a new 1.5 mL Eppendorf lobind tube. This was repeated twice to obtain 40 µL of extract. The final DNA concentration was measured using Qubit fluorometer 3.0 with HS reagents and used directly for downstream shotgun library preparation.

### 2. DNA Sequencing and Mapping

#### 2.1 Shotgun library preparation for Illumina

For the library preparation, 32 µL of extract was transferred to 0.5 mL Lobind Eppendorf tubes and mixed with an end-repair mastermix consisting of 0.4 µL T4 DNA polymerase (3 U/µL, NEB), 1 µL T4 polynucleotide kinase (10 U/µL, NEB), 0.4 µL dNTP (25 mM stock), 4 µL T4 DNA ligase reaction buffer (NEB) and 2.2 µL reaction buffer (25% PEG-4000, 2 mg/ml BSA, 400 mM NaCl).

The end-repair reaction was incubated in a thermal cycler for 30 minutes at 20°C, followed by 30 minutes at 65°C and cooled to 4°C. Two µL of blunt end Illumina adapter (as used in (Mak et al. 2017)) 10 µM stock was added and the sample was mixed by flicking. A ligation mastermix was subsequently added, consisting of: 1 µL T4 DNA ligase reaction buffer (NEB), 6 µL PEG-4000 (50%) and 1 µL T4 DNA ligase (400 U/µL, NEB). The ligation reaction was incubated at 20°C in a thermocycler for 30 minutes followed by 10 minutes at 65°C and cooled to 4°C. The sample was then added to a Fill-in mastermix consisting of 2 µL isothermal amplification buffer (NEB), 0.8 dNTP (25mM stock), 1.6 µL Bst 2.0 Warmstart polymerase (NEB) and 5.6 µL molecular grade water. The sample was incubated for 15 minutes at 65°C, followed by 15 minutes at 80°C and cooled to 4°C. Finally, the libraries were purified using 100 µL SPRI bead solution (Rohland and Reich 2012).

#### 2.2 Quantitative PCR (qPCR)

To estimate the number of library molecules produced, a qPCR was performed using Roche Lightcycler 480 SYBR green I reagents (Thermo Fischer scientific). Samples were diluted 1:100 in EBT buffer and 4.75 µL was used in a 10 µL reaction with 5 µL Roche Lightcycler mastermix and 0.25 µL IS7/8 primer mix (10 µM) (Meyer and Kircher 2010). The reactions were run on an Agilent mx3005 instrument. Based on the amplification seen in qPCR, it was estimated to give the libraries 15 cycles in indexing PCR.

#### 2.3 Index PCR amplification

Index amplification was performed with index primers, with newly designed indexes to achieve matching dual seven-base indices. Thus, the overall design of the primer was similar to the setup described in Kircher et al. (2012), but with newly designed matching indices of seven bases instead of six. This ensured a minimum three-base difference between any two pairs of indices from the list of 12 we used. Primers were ordered from IDT, LubioScience GmbH (https://eu.idtdna.com/pages) and prepared as described in Kircher et al. 2012.

Samples were amplified using Kapa Biosystems Kapa HIFI U+ PCR mastermix, in a setup with 23 µL undiluted sample, 1 µL of each primer (10 µM) and 25 µL Kapa HIFI U+ PCR mastermix. Incubation was performed for two minutes at 94 °C followed by

15 cycles of 20 seconds at 94°C, 30 seconds at 60°C, one minute at 72°C and finally two minutes at 72 °C. PCR products were purified using SPRI beads, as previously described, using 60 μL SPRI bead solution. PCR products were quantified and analyzed on an Agilent 2100 Bioanalyzer prior to pooling and sequencing.

Shotgun sequencing was performed at the Danish National high-throughput sequencing centre, Copenhagen Denmark, on one lane on an Illumina Hiseq 2500 instrument in paired end mode, running 125 cycles.

#### 2.4 Genome Mapping

Sequenced libraries were examined with the software FastQC (https://www.bioinformatics.babraham.ac.uk/projects/fastqc/). Reads were then processed to remove sequencing adapters using cutadapt 1.9.2 (Martin 2011). Clipped reads were mapped against the Sal1, 3D7 and PvP01 reference genomes using BWA aln (Li and Durbin 2009), with no trimming, disabling edit distance and setting a gap penalty opening of two. Alignments were generated with paired-end reads using BWA sampe. Reads exhibiting mapping qualities below 30 were removed using picard 2.8.3 (https://broadinstitute.github.io/picard/). Duplicated reads were removed using Samtools 1.6 (Li et al. 2009). Mapped unique reads were examined for postmortem damange with mapDamage2 (**Supplementary Figure 1**) (Jónsson et al. 2013) to determine the authenticity and the degradation rate of the data. Qualimap 2 (Okonechnikov et al. 2015) was used to obtain quality and coverage statistics from all mapped data.

### 3. Human contamination

Sequenced reads were mapped against the Human reference genome (GRCh37) (GCA_000001405.1) following the same strategy as described for the *P. vivax* mapping. The number of human reads was used to determine both the efficiency of the human Capture depletion, and to determine the rate of *Plasmodium/Human* sequenced and mapped reads. A total of 83,229,879 sequenced reads mapped to the human genome. The resultant ratio of Plasmodium/human mapped reads was 0.89%. However, 90,864,016 sequenced reads did not map to either the human genome or to the *P. vivax / P. falciparum* genomes.

In order to study the origin of the unmapped reads, we subsampled one million reads out of the 90,864,168 unmapped reads that did not map to *P. vivax, P. falciparum* or *H. sapiens.* We used BLAST to determine the origin of these reads: 2658 out of the one million reads could be assigned to a BLAST hit with 2639 of them mapping to complex regions of the human genome assembly, whilst the rest were unspecific hits.

### 4. Allele and haplotype sharing analyses

To better characterize fine-scale genetic structure amongst strains included in the population genetics dataset (**Supplementary Table 10**), we inferred patterns of allele and haplotype sharing using CHROMOPAINTER v2 (Lawson et al. 2012). We first subset the population genetics dataset to only sites present in Ebro-1944 and then retained only those samples with ≤10% missing data (77,420 sites, 218 samples).

We initially implemented CHROMOPAINTER using an unlinked approach (*-u* switch) which considers variant sites as fully independent and thus does not rely on our ability to confidently call haplotypes. Under the unlinked model CHROMOPAINTER calculates, separately for each position, the probability that a "recipient" chromosome is most closely related to a particular "donor" in the dataset. Here, we use all strains in our reduced global dataset as donors and the equivalent strains as recipients in an "all-versus-all" painting approach so that for each recipient sample *r*, we define $y_d^r$ to be the total amount (measured in matching allele counts) of DNA for which recipient sample *r* is inferred to be most closely related to a donor chromosome from group *d* (the "chunkcounts.out" output from the CHROMOPAINTER model). We clustered the resulting allele-sharing profiles in fineSTRUCTURE (Lawson et al. 2012), grouping strains based on each of their inferred $y_d^r$ in relation to every other sample. fineSTRUCTURE was run sampling 2,000,000 MCMC iterations (-y) with a thinning interval of 10,000 (-z). The tree building step, after discarding the burn-in, was applied for 100,000 hill climbing iterations (-x) and 10,000 tree comparisons (-t) (**Supplementary Figure 8**).

We repeated our unlinked chromosome painting analyses using a linked CHROMOPAINTER model, which now considers the correlation among neighbouring sites (e.g. haplotype information). As the linked CHROMOPAINTER model currently relies on no missing sites, we performed imputation on the 10% of sites that were not covered in our dataset of 77,420 sites and 218 samples. We followed the imputation protocol set out by Samad et al (2015) for *P. falciparum* using BEAGLE v3.3.2 (Browning and Browning 2013). Using this imputed dataset, we then repeated our chromosome painting procedure. However, rather than calculating the probability at each position independently, we now consider linkage assuming a uniform recombination map of constant rate 1cM every 14kb. Under the linked model, $y_d^r$ now defines the length or proportion of total DNA shared between each recipient sample *r* and each donor *d* (the "chunklengths.out" output from linked CHROMOPAINTER v2 model). We first inferred the switch rate ($N_e$) and mis-copying parameter (θ) as advocated by Lawson et al. (2012) by running the CHROMOPAINTER expectation-maximisation (E-M) algorithm on every recipient strain for 10 iterations. The inferred values were weight-averaged across all chromosomes to give mean estimates of $N_e$=1843.318 and θ=0.003 across the 218 samples. CHROMOPAINTER was then run with these values fixed using the -N and -M switches.

We found a strong correlation between our inferred painting profiles under an unlinked and linked CHROMOPAINTER model ($r^2$=0.94) suggesting our imputation procedure provides robust calls and consistent demographic inference (**Supplementary Figure 10**). Thus, as before, we performed clustering on the inferred painting profiles across all samples. We ran fineSTRUCTURE using an estimated normalisation parameter, *c*=0.014. fineSTRUCTURE's inferred hierarchical clustering under these separate implementations are provided in **Supplementary Figures 8-9**.

In order to include more strains from diverse populations, we extracted one further subset from our original global population genetics dataset collection but now allowing for samples to have ≤30% of sites missing. This allowed the incorporation of an additional 100 strains from geographically relevant regions. As before, we followed the protocol of Samad et al. (2015) to impute missing sites using BEAGLE v3.3.2 (Browning and Browning 2013). We again estimated the CHROMOPAINTER v2 switch and mis-copying parameters, providing estimates of $N_e$=1261.830 and

$\theta$=0.003 which were fixed and run in CHROMOPAINTER and fineSTRUCTURE (*c*=0.009). We assessed the reliability of our imputation and painting procedure by comparing our inferred coancestry matrices to those inferred without imputation (unlinked model), and following imputation (10% and 30% respectively) (**Supplementary Figure 10**). fineSTRUCTURE's inferred hierarchical clustering following imputation of samples with up to 30% of sites missing is provided in **Figure 3a**.

### SUPPLEMENTARY FIGURES

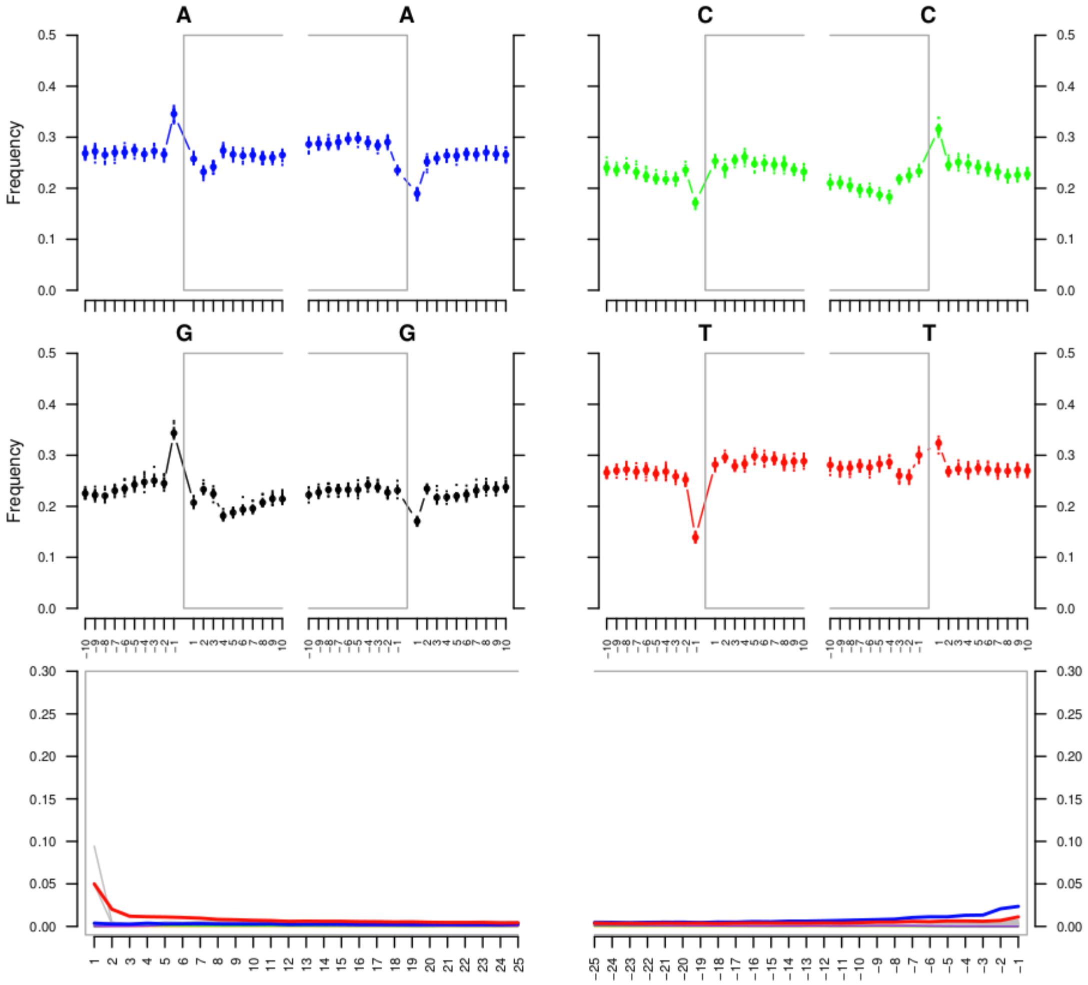


**Supplementary Figure 1:** MapDamage2 mis-incorporation plot to visualize patterns of post-mortem damage. MapDamage2 was applied to the data extracted and sequenced in 2017. The top two rows provide the base frequency of A, C, G, T respectively outside and inside (grey box) the reads. The bottom row provides the position specific substitutions from the 5' (left) to 3' (right) (x-axis) with red providing the C to T substitution frequency, blue providing the G to A substitution frequency and grey providing the frequency of all other substitutions.

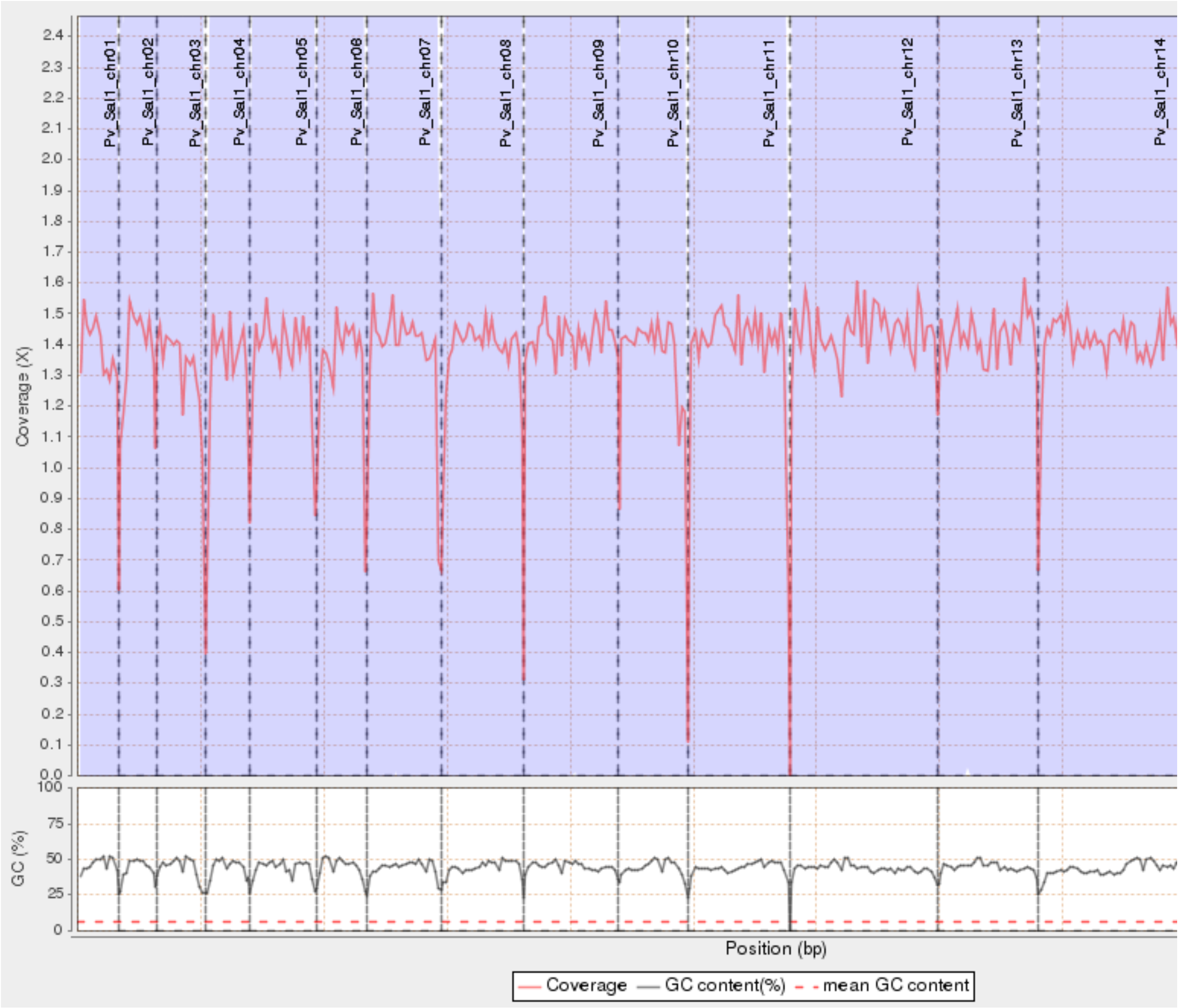


**Supplementary Figure 2:** Top: Coverage distribution of the Ebro-1944 sample along the *P. vivax* Sal1 reference chromosomes plotted with Qualimap 2. Bottom: GC content (%) distribution of Ebro-1944 along the chromosomes of the Sal-1 *P. vivax* reference genome.

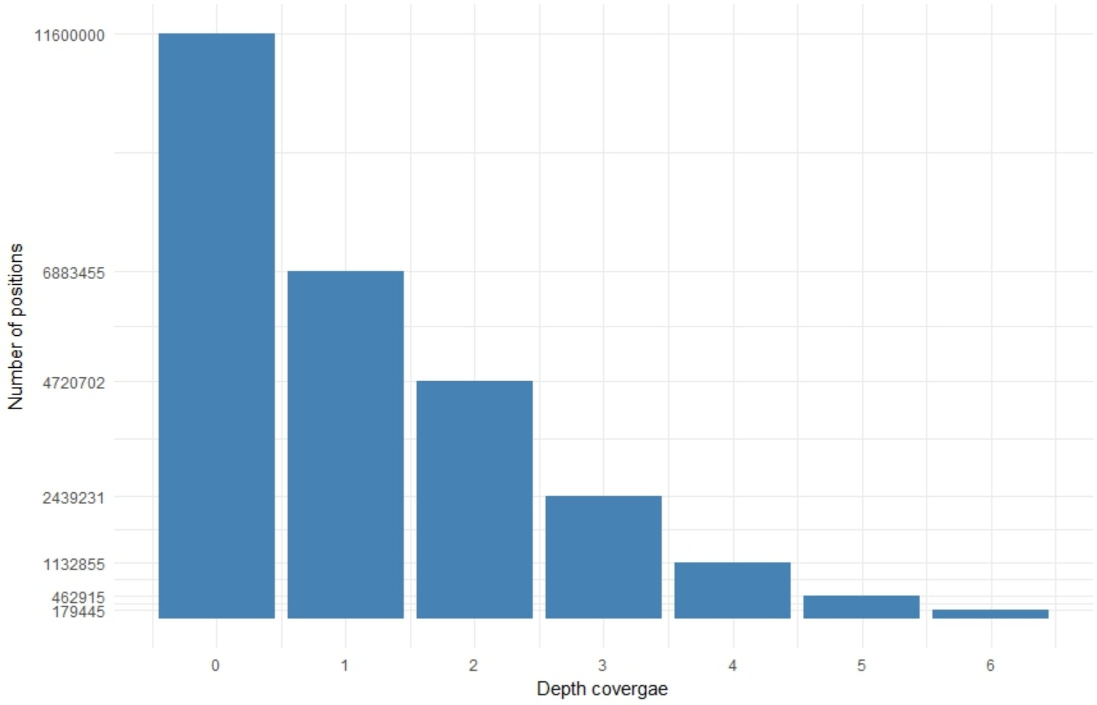


**Supplementary Figure 3:** Histogram of the coverage per site distribution of the Ebro-1944 sample mapped to the *P. vivax* Sal1 reference genome.

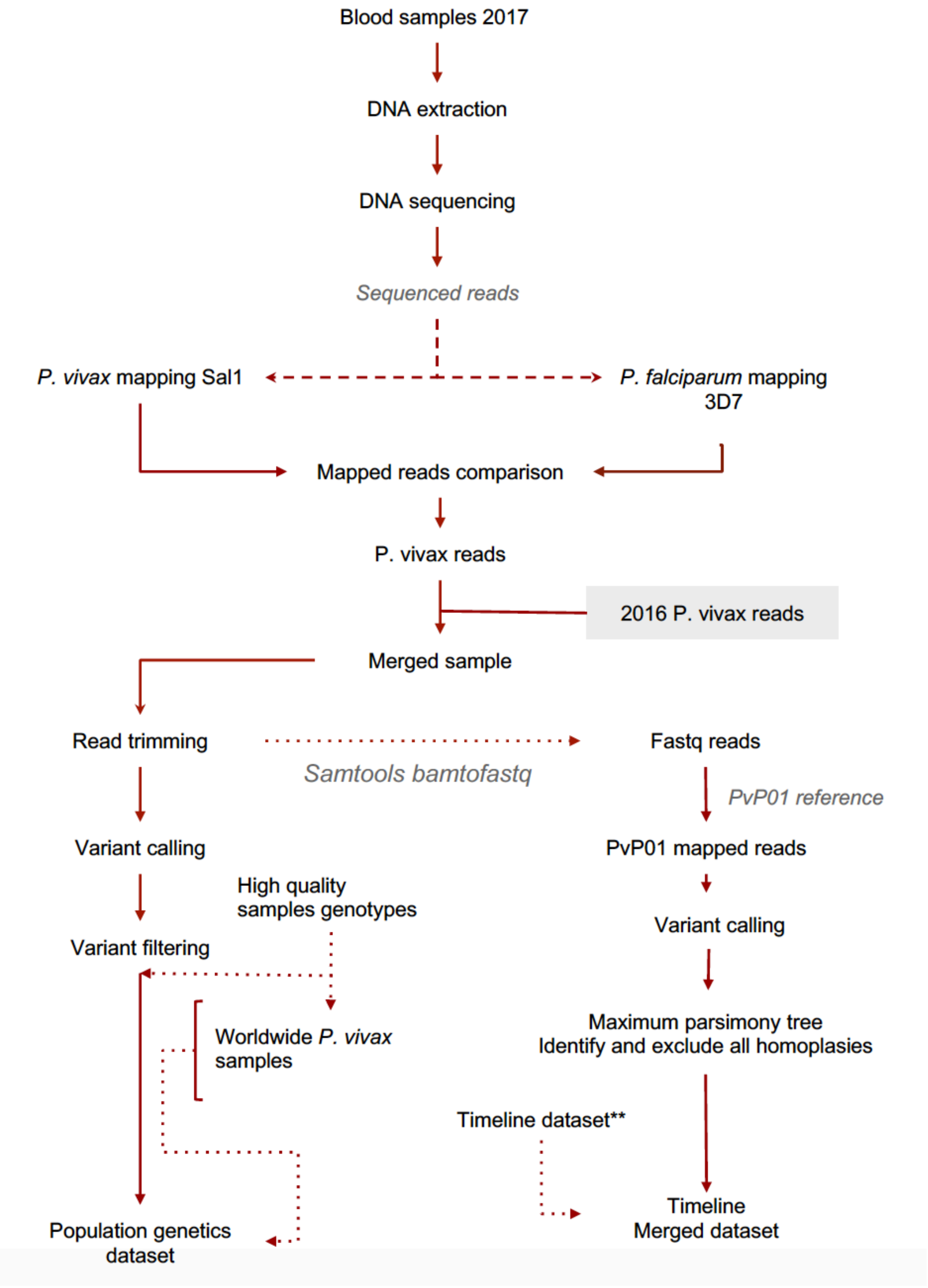


**Supplementary Figure 4:** Flux diagram (I) providing steps in the processing of the Ebro-1944 genome and in the dataset preparation for the population genetics dataset and the timeline phylogenetics dataset.

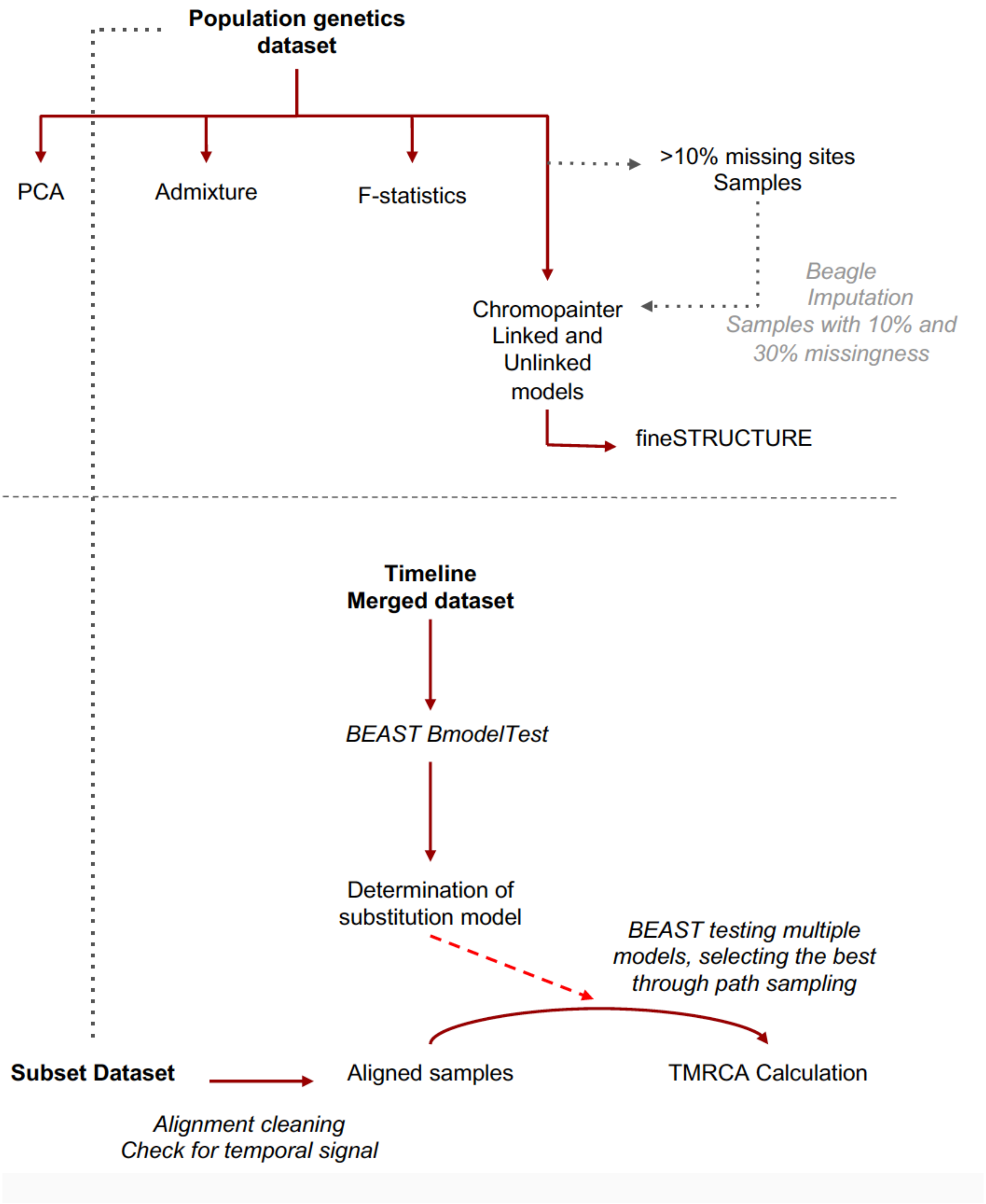


**Supplementary Figure 5:** Flux diagram (II) detailing the analyses performed on the population genetics dataset and the timeline phylogenetic dataset.

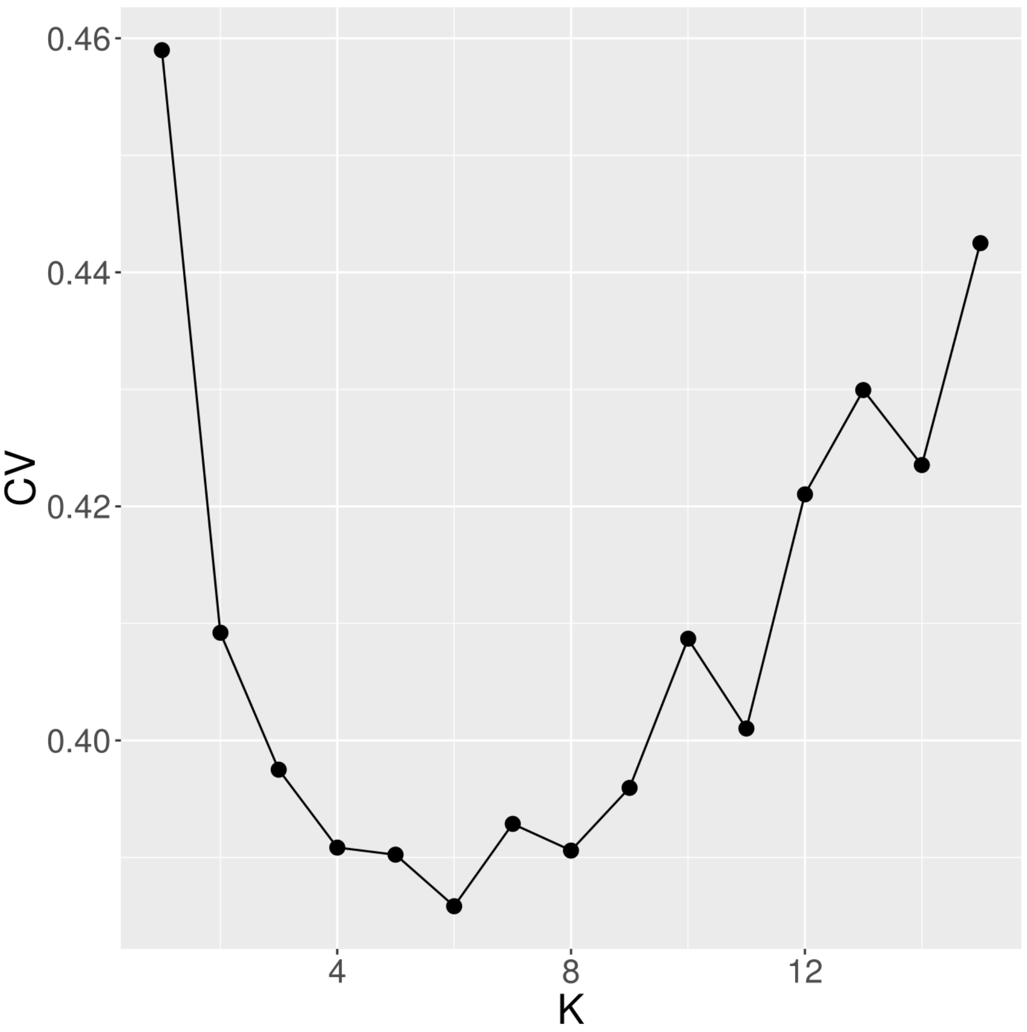


**Supplementary Figure 6:** Cross-validation (CV) scores (y-axis) following ADMIXTURE's default five-fold cross-validation procedure. CV scores were generated from unsupervised ADMIXTURE analyses applied to the population genetics dataset over 15 values of K (x-axis). The lowest cross-validation error was obtained at K=6, with the admixture profile at K=6 provided in main text **Figure 1c**.

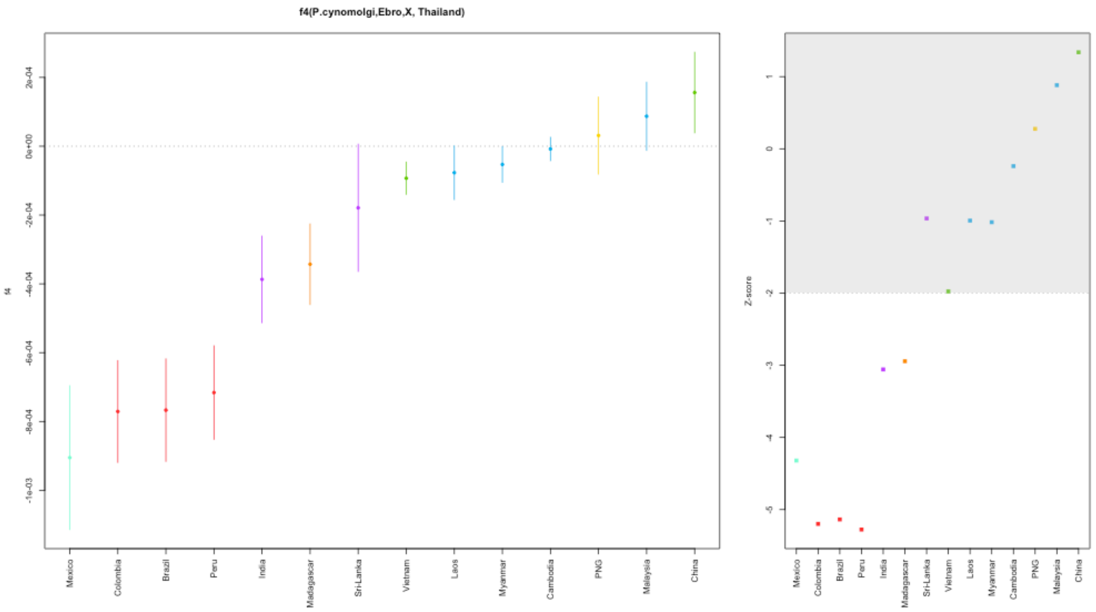


$f_4$(*P. cynomolgi,* Ebro-1944; X, Thailand)

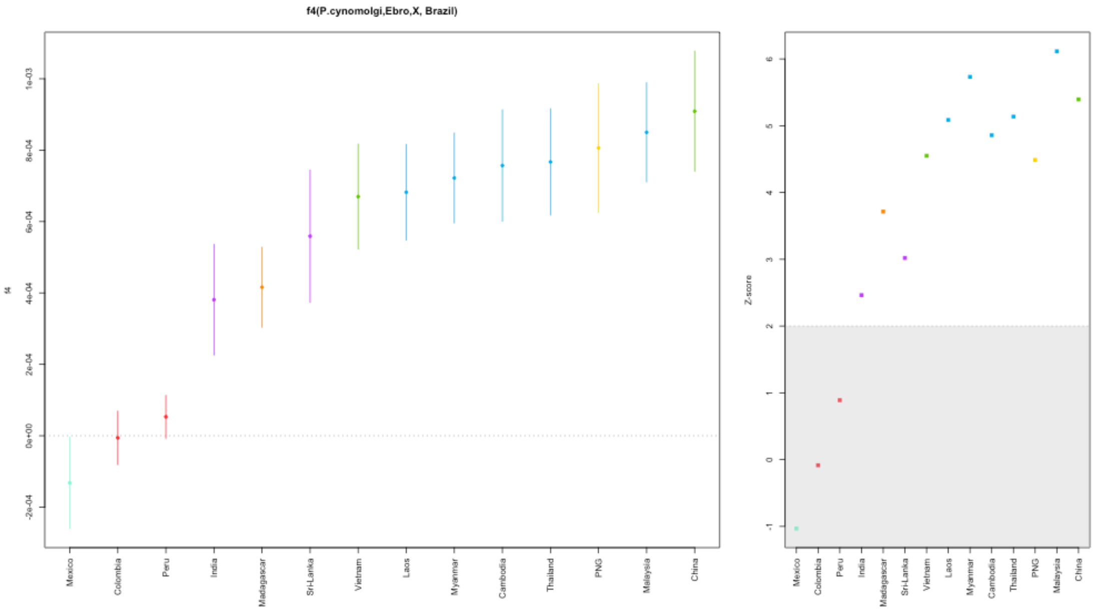


$f_4$(*P. cynomolgi,* Ebro-1944; X, Brazil)

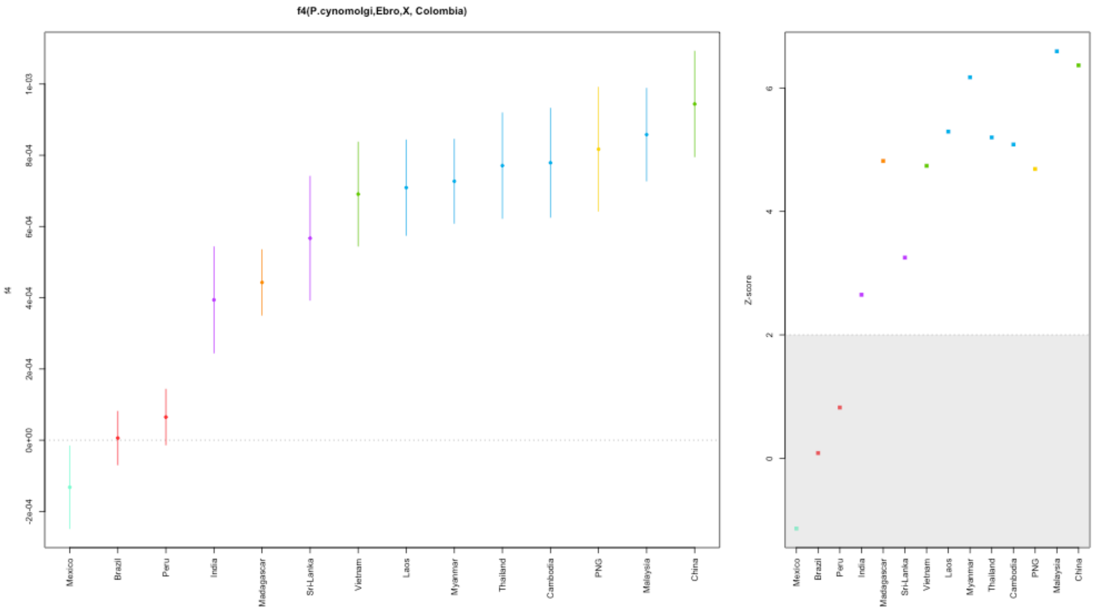


$f_4$*(P. cynomolgi,* Ebro-1944; X, Colombia)

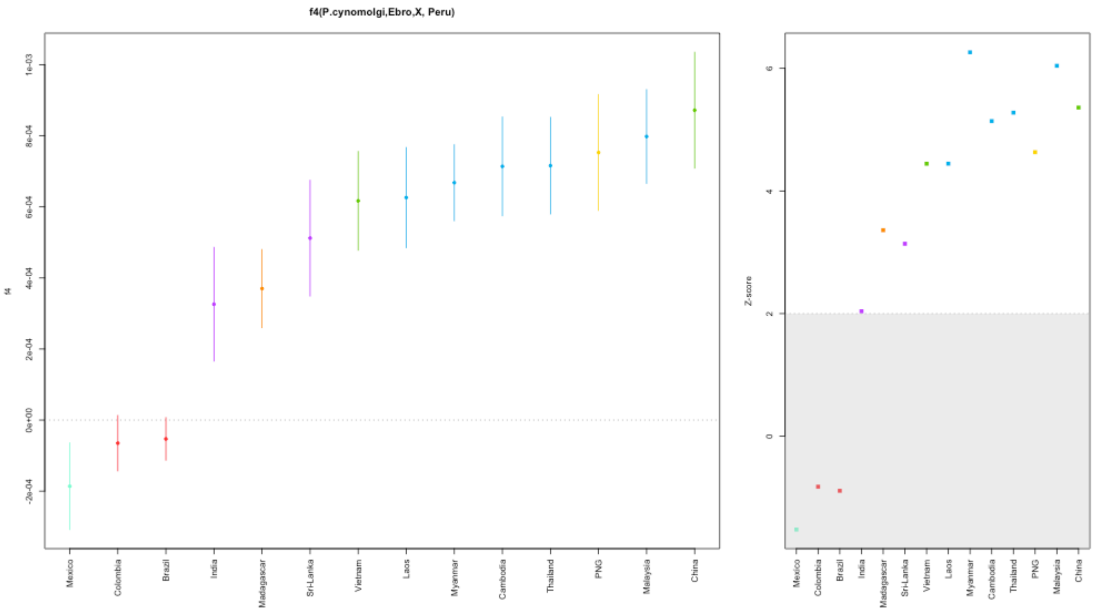


$f_4$*(P. cynomolgi,* Ebro-1944; X, Peru)

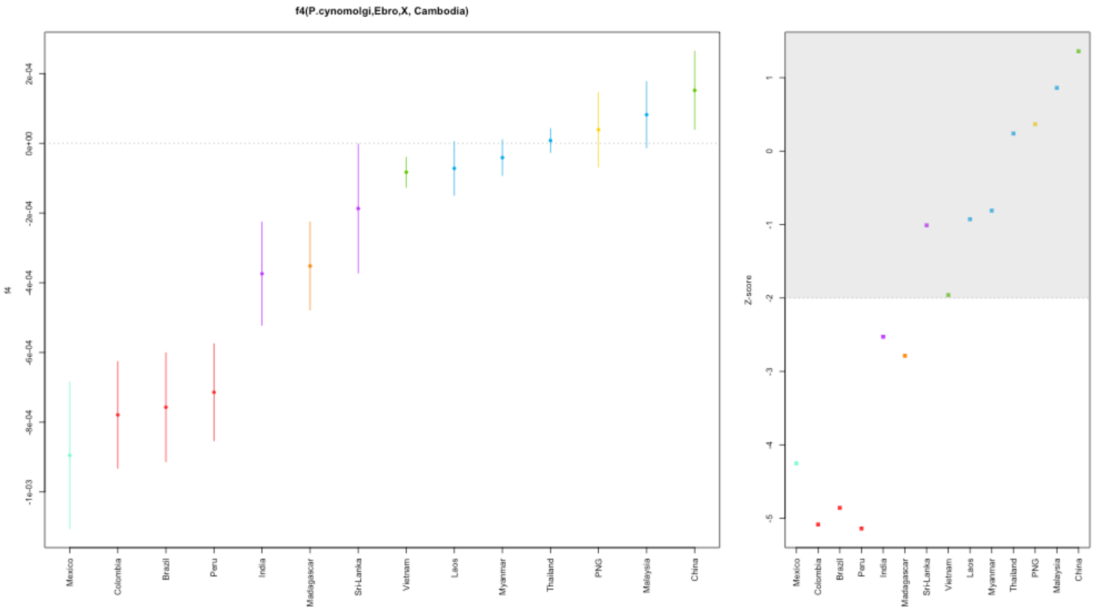


$f_4$(*P. cynomolgi,* Ebro-1944; X, Cambodia)

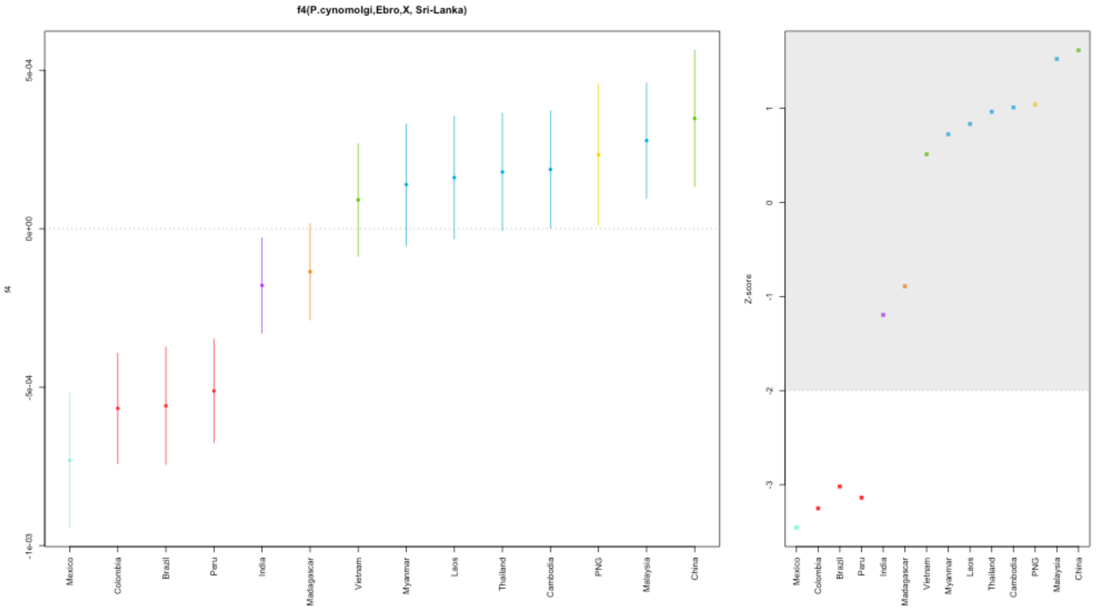


$f_4$(*P. cynomolgi,* Ebro-1944; X, Sri-Lanka)

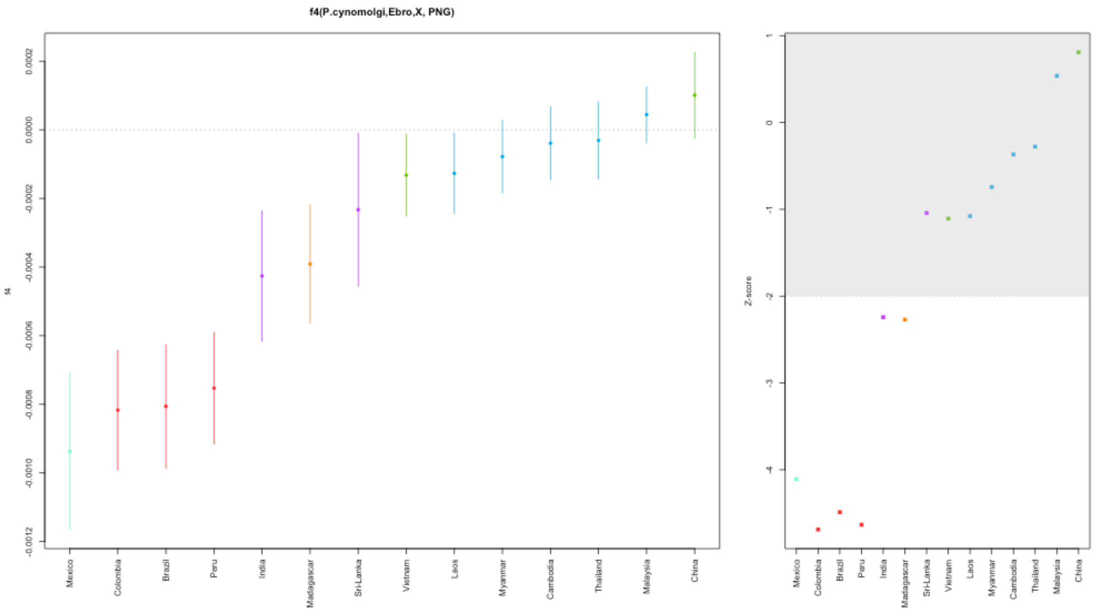


*f*4(*P. cynomolgi,* Ebro-1944; X, Papua-New-Guinea)

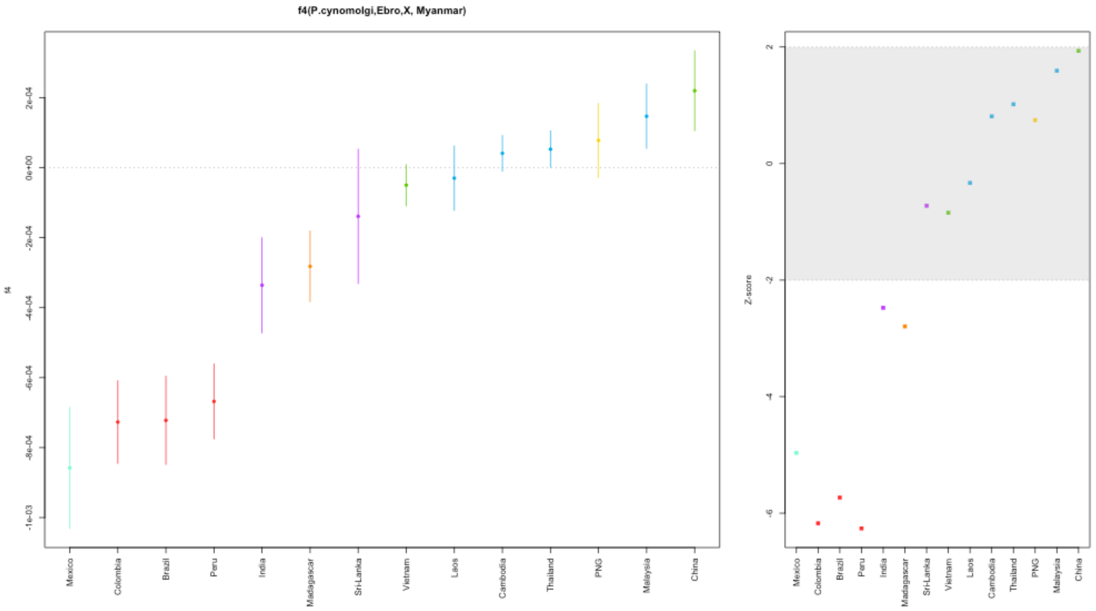


*f*4*(P. cynomolgi,* Ebro-1944; X, Myanmar)

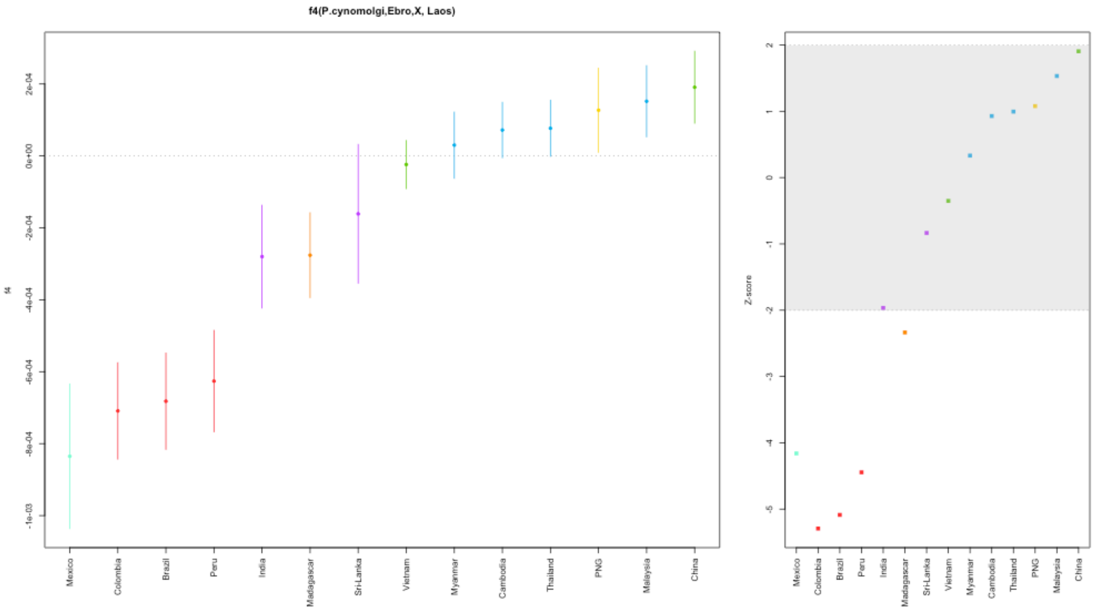


$f_4$*(P. cynomolgi,* Ebro-1944; X, Laos)

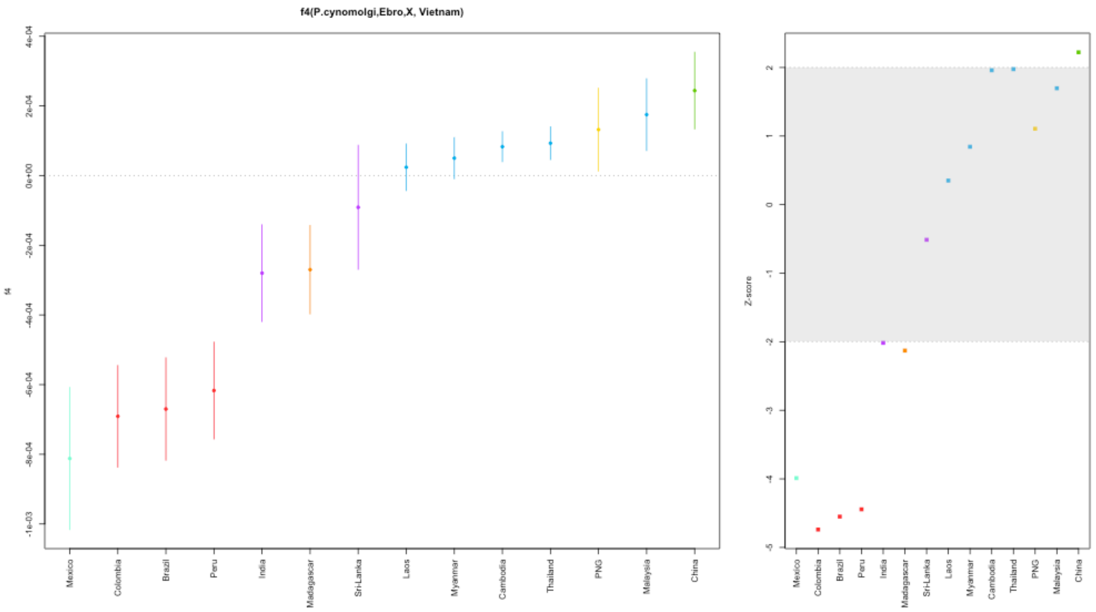


$f_4$*(P. cynomolgi,* Ebro-1944; X, Vietnam)

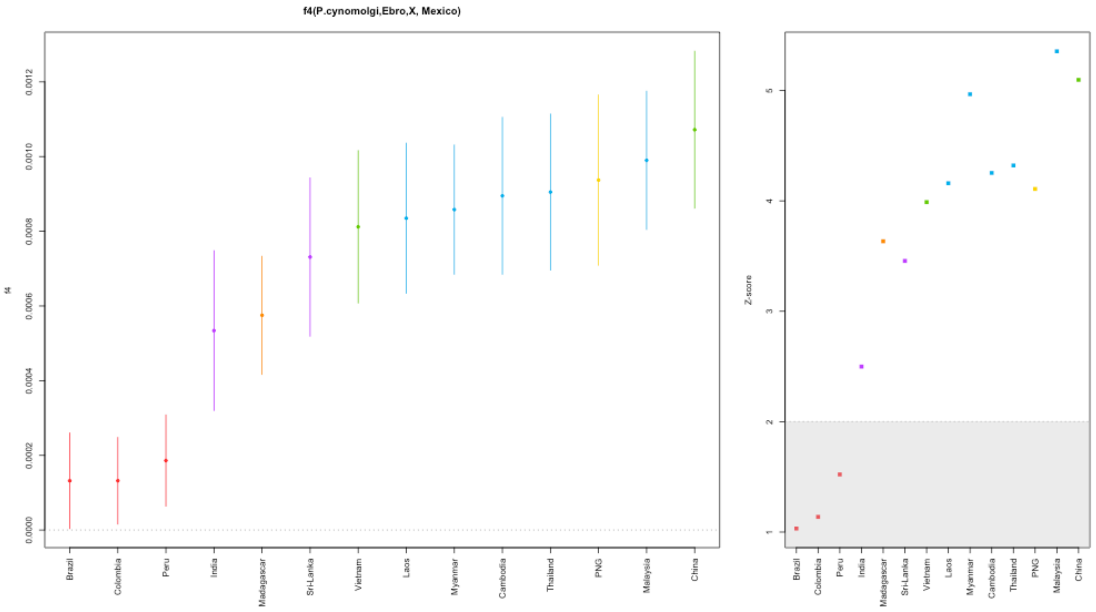


*$f_4$(P. cynomolgi,* Ebro-1944; X, Mexico)

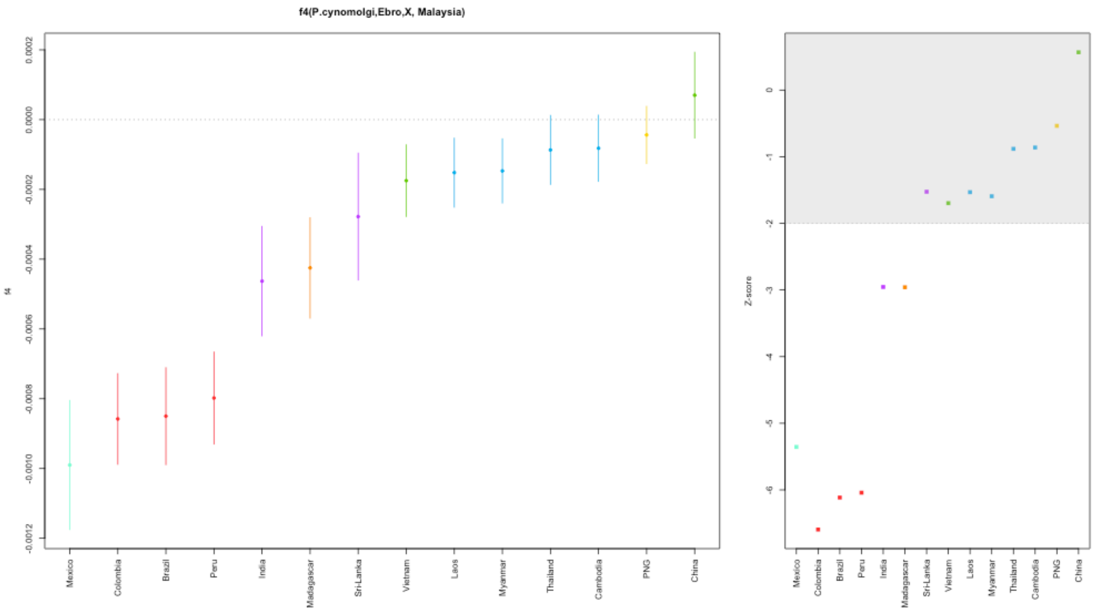


*$f_4$(P. cynomolgi,* Ebro-1944; X, Malaysia)

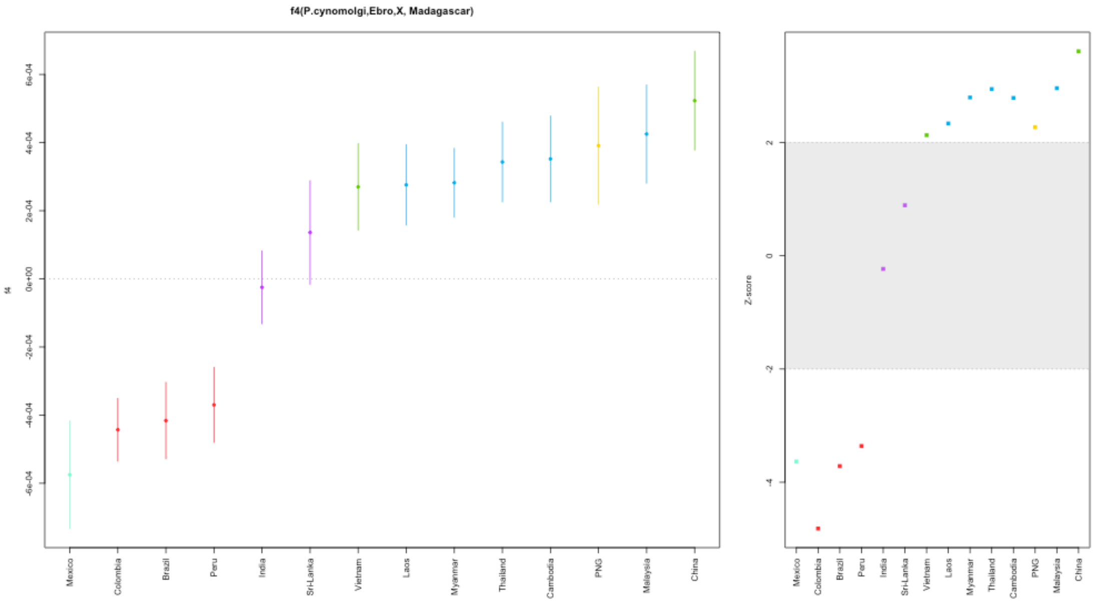


$f_4$*(P. cynomolgi,* Ebro-1944; X, Madagascar)

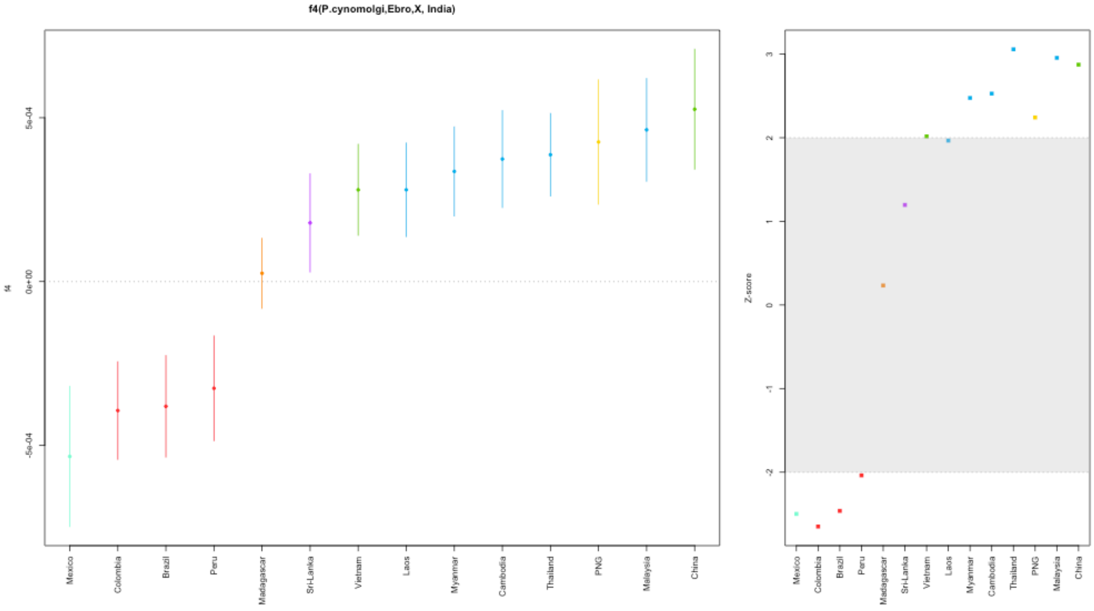


$f_4$*(P. cynomolgi,* Ebro-1944; X, India)

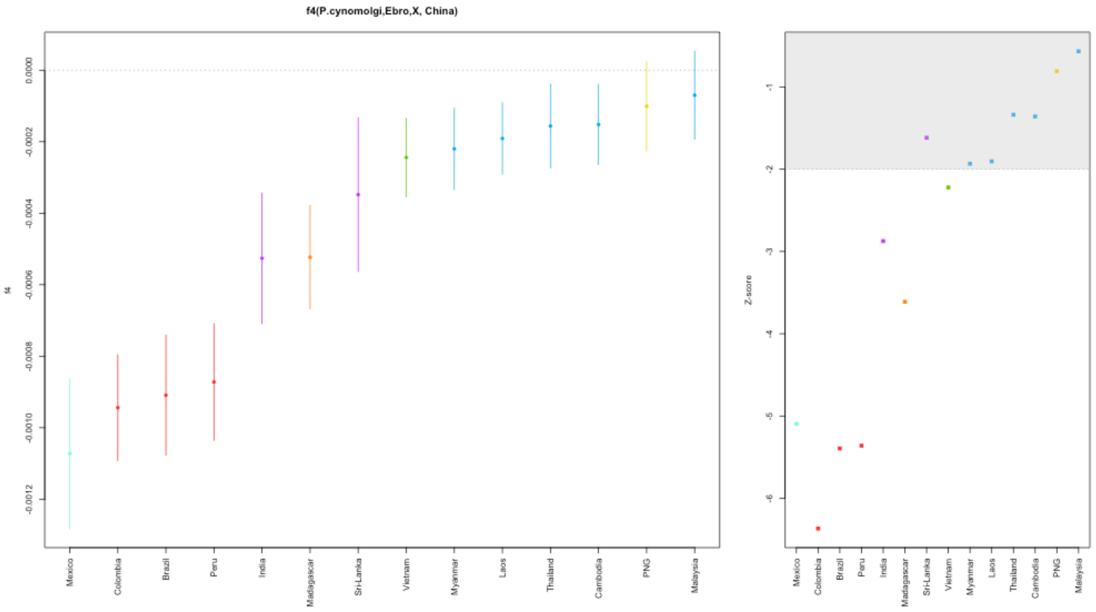


*f*4(*P. cynomolgi,* Ebro-1944; X, China)

**Supplementary Figure 7:** *f*4 population test statistics, plus standard errors, for combinations of populations included in the population genetics dataset. The selected populations are listed below each plot, coloured as in main text **Figure 1a**. Z scores are provided in the right panel of each plot with significant absolute Z scores >2 falling outside of the grey shaded box.

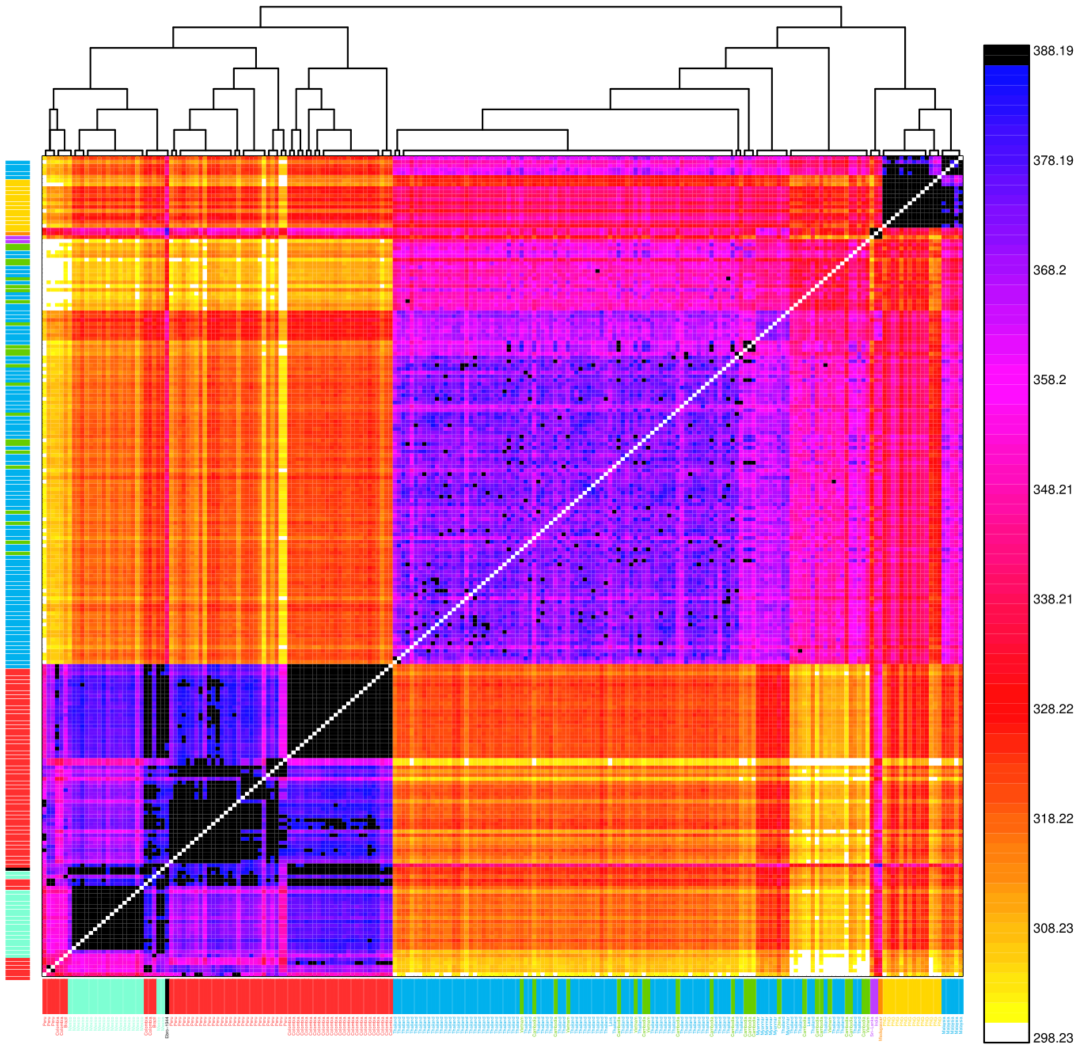


**Supplementary Figure 8:** CHROMOPAINTER's inferred counts of matching DNA between all included *P. vivax* samples. Each of the 27 inferred clusters (columns) is painted by each of the 27 clusters (rows). The tree at top shows fineSTRUCTURE's inferred hierarchical merging of these 27 clusters and the colours on the axes give the continental region and population to which samples in each cluster are assigned. Ebro-1944 is depicted in black and clusters with 2 samples from Mexico, 1 sample from Brazil and 2 samples from Colombia.

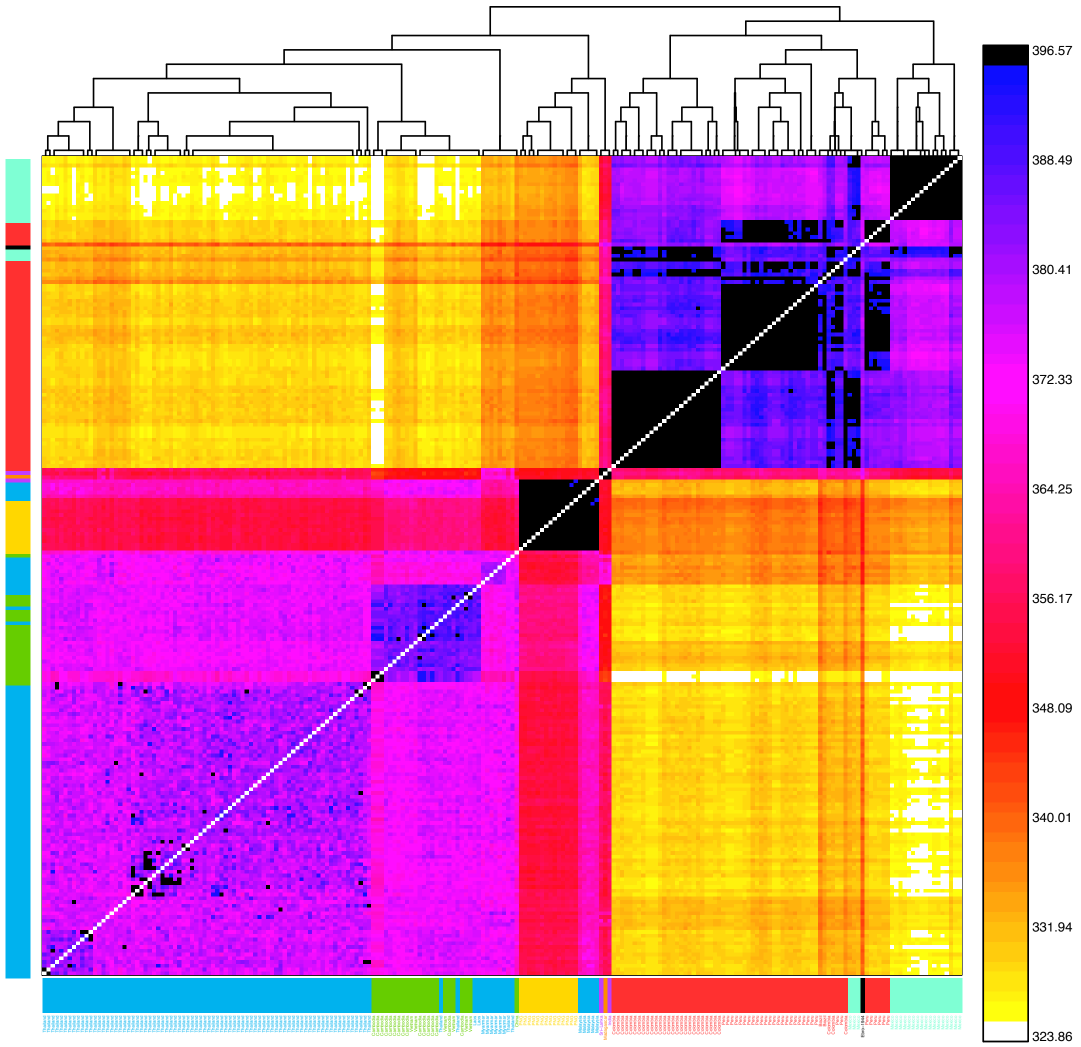


**Supplementary Figure 9:** CHROMOPAINTER's inferred counts of matching DNA between all included *P. vivax* samples. Each of the 70 inferred clusters (columns) is painted by each of the 70 clusters (rows) using a linked model following imputation of 10% of sites. This figure is comparable to **Supplementary Figure 8** once taking linkage i.e. haplotype information into account. The tree at top shows fineSTRUCTURE's inferred hierarchical merging of these 70 clusters and the colours on the axes give the continental region and population to which samples in each cluster are assigned. Ebro-1944 is depicted in black and clusters with samples from Mexico and Peru.

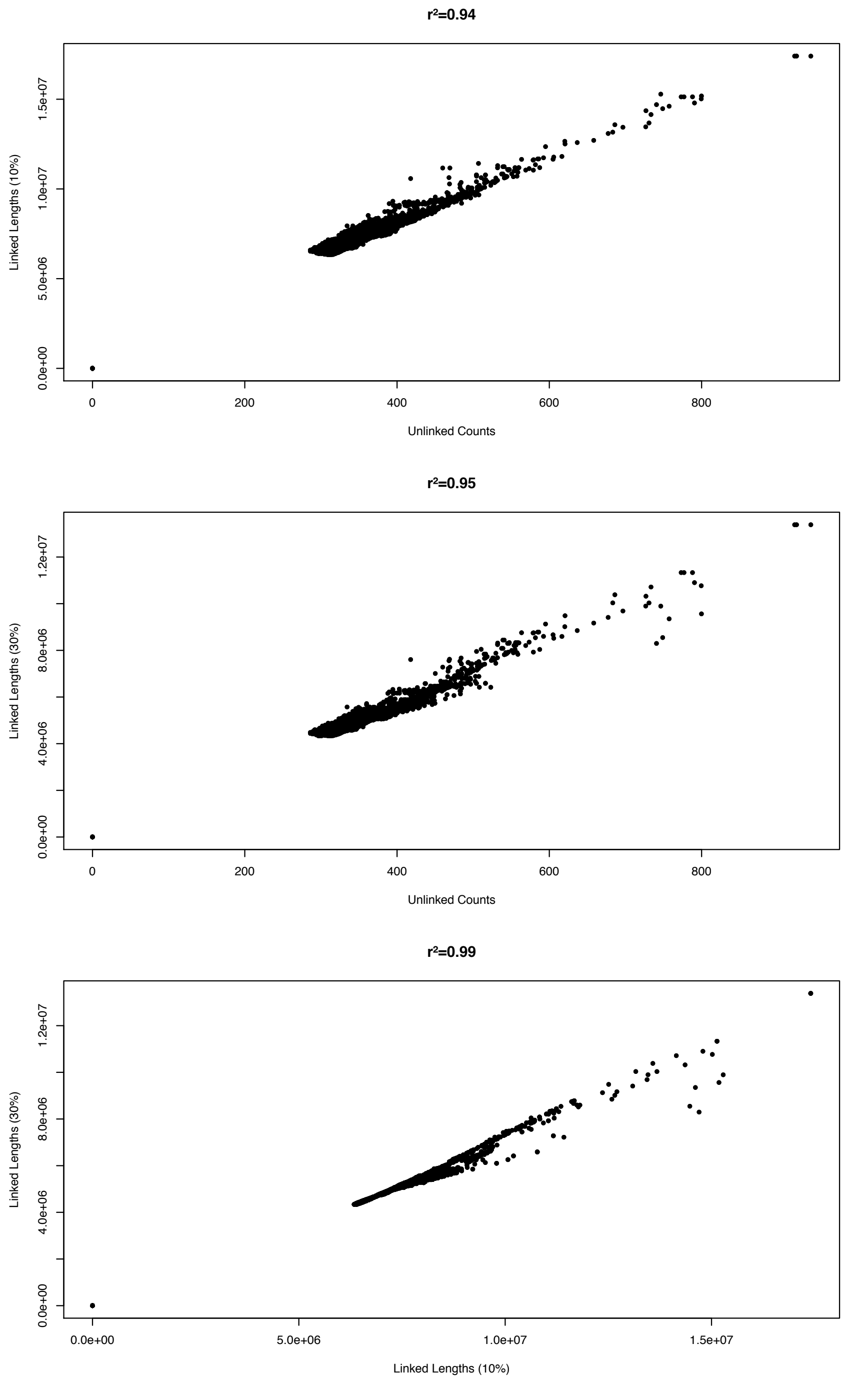


**Supplementary Figure 10:** Correlations between CHROMOPAINTER inferred chunkcounts (unlinked) and chunklengths (linked) across three independent chromosome painting analyses: Unlinked using data with up to 10% missingness across 218 samples, Linked (10%) using data with sites with up to 10% missingness imputed using BEAGLE across 218 samples, Linked (30%) using data with sites with up to 30% missingness imputed using BEAGLE across 318 samples, see main text Methods and **Supplementary Section 4**.

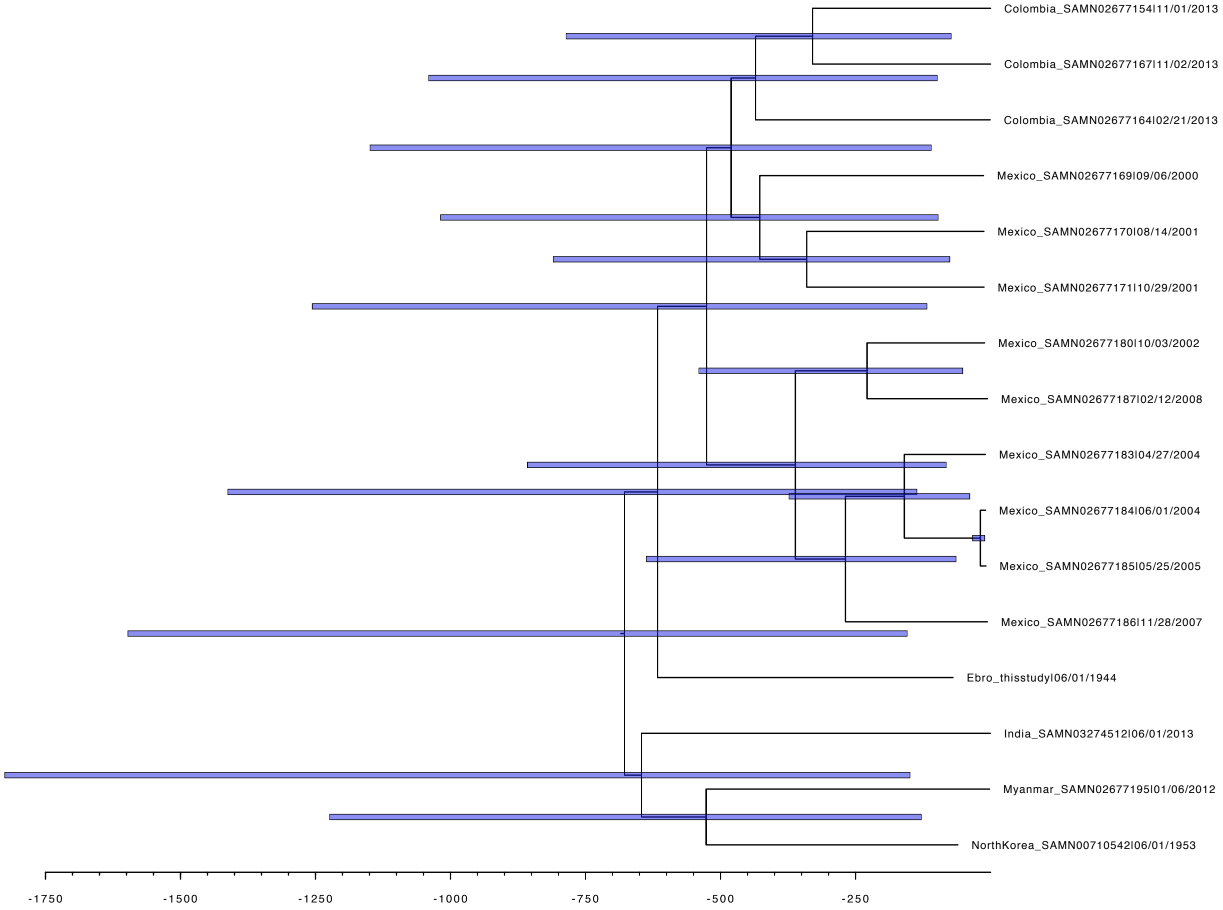


**Supplementary Figure 11** Maximum clade credibility tree having discarded a 10% burn-in under a Coalescent Bayesian Skyline implementation of BEAST 2 (Bouckaert et al. 2014) with a relaxed molecular clock. The x-axis provides the dated phylogeny in years before 2013. Blue bars provide the 95% HPD estimates around each node. Full results for different demographic models and clock rate priors are provided in **Supplementary Table 5**.

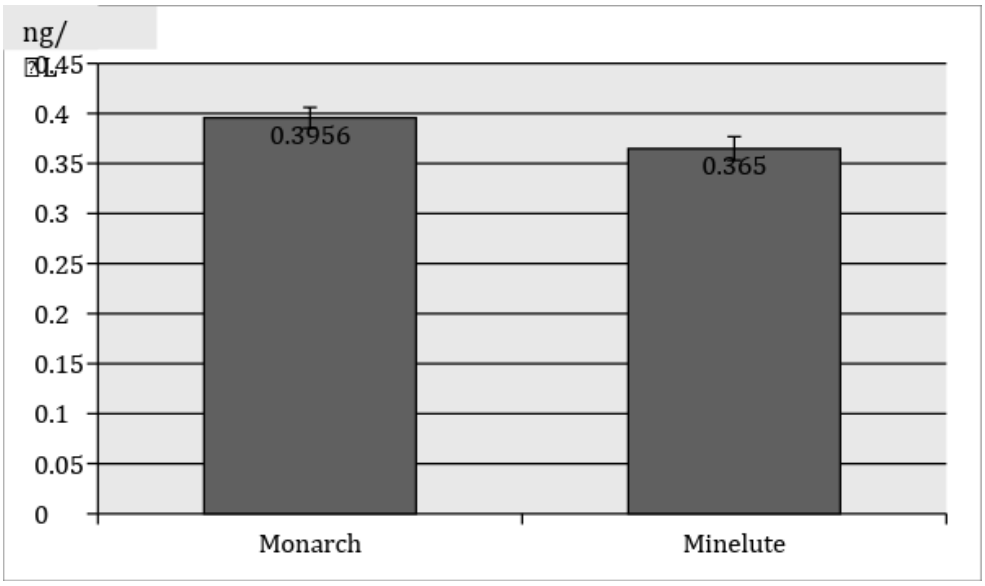


**Supplementary Figure 12**: 50 bp PCR fragments were dissolved in EB buffer and added 10x volume of binding buffer. Then passed through the columns at 3000xg, washed with 750 µL 80% ethanol at 10.000xg for one minute and dry spun at 17.000xg, before adding 20 µL of EB buffer, incubating at 37°C for 10 minutes before collecting. This was done twice and the eluate measured using a Qubit 2.0 fluorometer, using HS reagents and 10 µL input for precision. Error bars provide the standard deviation (*n*=5).

### SUPPLEMENTARY TABLES

**Supplementary Table 1:** Sequence reads and mapping coverage statistics. The mapping results are presented for the data extracted in 2016 and 2017 and for the combined *P. vivax* reads used to generate the Ebro-1944 genome.

| Sample | Sequenced reads | Mapped reads | | Number of mapped reads after duplicate removal (rmdup) | | Number of mapped reads after quality removal (Q30) | |
|---|---|---|---|---|---|---|---|
| | | Genome | Mitochondria | Genome | Mitochondria | Genome | Mitochondria |
| *P. vivax* 2016 | 468134047 | 687725 | 1052590 | 75861 | 1796 | 52643 | 459 |
| *P. vivax* 2017 | 255069630 | 742061 | 3784 | 432018 | 1964 | 428631 | 1962 |
| Ebro-1944 | - | - | - | - | - | 481245 | 2421 |

| *Table 1 continued* | Coverage estimate after Q>30 and duplicate removal | | % of covered genome | |
|---|---|---|---|---|
| Sample | Genome (mappable) | Mitochondria | Genome | Mitochondria |
| *P. vivax* 2016 | 0.12 | 4.95 | - | - |
| *P. vivax* 2017 | 1.28 | 27.19 | - | - |
| Ebro-1944 | 1.40 | 32.23 | 66.42% | 100.00% |

**Supplementary Table 2:** The number and coverage of mapped reads to the *P. vivax* Sal1 reference genome. The table provides the number of mapped and processed reads, the number of recovered bases and the mean depth of coverage values in the chromosomes of the Ebro-1944 *P. vivax* genome.

| Chromosome | Mapped unique reads (Q30) | Base count | Mean depth of coverage |
|---|---|---|---|
| Pv_Sal1_chr01 | 830022 | 1138687 | 1.3718756 |
| Pv_Sal1_chr02 | 755035 | 1039540 | 1.3768103 |
| Pv_Sal1_chr03 | 1011127 | 1319170 | 1.3046531 |
| Pv_Sal1_chr04 | 876652 | 1173030 | 1.3380794 |
| Pv_Sal1_chr05 | 1370936 | 1871490 | 1.3651184 |
| Pv_Sal1_chr06 | 1033388 | 1378855 | 1.3343052 |
| Pv_Sal1_chr07 | 1497819 | 2083111 | 1.3907628 |
| Pv_Sal1_chr08 | 1678596 | 2316326 | 1.3799186 |
| Pv_Sal1_chr09 | 1923364 | 2740402 | 1.4247963 |
| Pv_Sal1_chr10 | 1419739 | 1901532 | 1.3393532 |
| Pv_Sal1_chr11 | 2067354 | 2879446 | 1.3928170 |
| Pv_Sal1_chr12 | 3004884 | 4320788 | 1.4379217 |
| Pv_Sal1_chr13 | 2031768 | 2903455 | 1.4290288 |
| Pv_Sal1_chr14 | 3120417 | 4349299 | 1.3938198 |

**Supplementary Table 3:** Dataset of 338 *P. vivax* compiled from the literature and covering 128,081 genome-wide sites, used in PCA and ADMIXTURE analyses. This dataset represents a subset of the population genetics dataset described in **Supplementary Table 10**.

| Continent/Region | Country | Number |
|---|---|---|
| SouthEastAsia | Thailand | 112 |
| | Malaysia | 6 |
| | Laos | 2 |
| | Myanmar | 9 |
| EastAsia | Cambodia | 29 |
| | Vietnam | 11 |
| | China | 1 |
| CentralSouthAsia | India | 3 |
| | Sri-Lanka | 1 |
| Oceania | Papua_New_Guinea (PNG) | 24 |
| Central_America | Mexico | 20 |
| South_America | Brazil | 12 |
| | Colombia | 31 |
| | Peru | 72 |
| Africa | Madagascar | 4 |
| Europe | Spain | 1 |
| Total Samples, 60% missing threshold | | 338 |

**Supplementary Table 4:** Samples used in the timeline dataset for phylogenetic dating.

| Name | BioSample | Year |
|---|---|---|
| **Ebro_thisstudy\|06/01/1944** | This Study | 1944 |
| **Colombia_SAMN02677154\|11/01/2013** | SAMN02677154 | 2013 |
| **Colombia_SAMN02677164\|02/21/2013** | SAMN02677164 | 2013 |
| **Colombia_SAMN02677167\|11/02/2013** | SAMN02677167 | 2013 |
| **India_SAMN03274512\|06/01/2013** | SAMN03274512 | 2013 |
| **Mexico_SAMN02677169\|09/06/2000** | SAMN02677169 | 2000 |
| **Mexico_SAMN02677170\|08/14/2001** | SAMN02677170 | 2001 |
| **Mexico_SAMN02677171\|10/29/2001** | SAMN02677171 | 2002 |
| **Mexico_SAMN02677180\|10/03/2002** | SAMN02677180 | 2004 |
| **Mexico_SAMN02677183\|04/27/2004** | SAMN02677183 | 2004 |
| **Mexico_SAMN02677184\|06/01/2004** | SAMN02677184 | 2004 |
| **Mexico_SAMN02677185\|05/25/2005** | SAMN02677185 | 2005 |
| **Mexico_SAMN02677186\|11/28/2007** | SAMN02677186 | 2007 |
| **Mexico_SAMN02677187\|02/12/2008** | SAMN02677187 | 2008 |
| **Myanmar_SAMN02677195\|01/06/2012** | SAMN02677195 | 2012 |
| **NorthKorea_SAMN00710542\|06/01/1953** | SAMN00710542 | 1953 |

**Supplementary Table 5** Bayesian phylogenetic tip-dating results testing three priors on the demographic model and allowing for either a strict or relaxed molecular clock, lognormally distributed. The inferred clock rates are expressed as substitutions per base pair per year over the tested alignment and the 95% higher posterior density (HPD) intervals are provided in parentheses. The best model fit was selected based on the raw median likelihood and the maximum likelihood (ML) estimate following path sampling.

| Demographic Model | Clock Model | Clock Rate (mean 95% HPD) | Median Likelihood | Europe America split time (mean 95% HPD) | Path Sample ML |
|---|---|---|---|---|---|
| Coalescent Constant | Strict | $5.37E^{-7}$ ($4.71E^{-7}$ - $6.02E^{-7}$) | -40128243.82 | 358 (318-402) | $-4.012837E^{+07}$ |
| | Relax | $7.09E^{-7}$ ($1.31E^{-7}$ - $1.27E^{-6}$) | -40128031.92 | 420 (129-777) | $-4.013625E^{+07}$ |
| Coalescent Exponential | Strict | $5.35E^{-7}$ ($4.70E^{-7}$ - $6.01E^{-7}$) | -40128243.77 | 359 (317-400) | $-4.012836E^{+07}$ |
| | Relax | $6.34E^{-7}$ ($1.63E^{-7}$ - $1.12E^{-6}$) | -40128031.85 | 382 (149-775) | $-4.012825E^{+07}$ |
| Coalescent Bayesian Skyline | Strict | $5.45E^{-7}$ ($4.71E^{-7}$ - $6.01E^{-7}$) | -40128243.71 | 360 (319-402) | $-4.012840E^{+07}$ |
| | Relax | $5.57E^{-7}$ ($2.75E^{-8}$ - $1.06E^{-6}$) | -40128031.79 | 598 (136-812) | $-4.012821E^{+07}$ |

**Supplementary Table 6** (see extended excel file)
Annotation of the 4,800 SNPs analyzed with SNPeff. SNPs located in each gene have been classified by their effect.

**Supplementary Table 7:** Derived and new mutations detected in Ebro-1944 in genes related to drug resistance and host infectivity.

| **Gene** | **Gene function** | **Position** | **Mutation** | **Described** |
|---|---|---|---|---|
| *PvDHPS* (PVX_123230) | Drug Resistance | 14:1258389 | Met205Ile | (Hawkins et al. 2009) |
| *CLAG* (PVX_094265) | Cytoadherence-linked asexual gene | 08:72119 | Leu623Arg | This study |
| *PvMRP1* (PVX_097025) | Drug Resistance | 02:154391 | Val1478Ile | (Dharia et al. 2010) |
| *PvMRP1* (PVX_097025) | Drug Resistance | 02:158047 | Thr259Arg | (Dharia et al. 2010; de Oliveira et al. 2017) |

**Supplementary Table 8:** Allelic state of genetic variants previously associated with resistance to antimalaria drugs in the Ebro-1944 strain. The historical European strain shows the derived allele in only one position (M205I), of unknown effect, in a gene associated with resistance to Sulfadoxine.

| Gene | Position | Base Change | AA Change | Known Effect | Ebro-1944 Genotype | Reference |
|---|---|---|---|---|---|---|
| Chloroquine resistance transporter (PvCRT) | Chr01:330,981 | A -> AAAG | Insertion | Chloroquine Resistance | Not Analyzed | (Suwanarusk et al. 2007) |
| Multidrug Resistance (MDR) | Chr09:362,870 | A → G | F1076L | Antimalarial multidrug resistance | Reference variant | (Brega et al. 2005) |
| | Chr09:363,169 | T → A | Y958F | Antimalarial multidrug resistance | Reference variant | (Brega et al. 2005) |
| | Chr09:363,223 | G → A | T958M | Antimalarial multidrug resistance | Not present | (Sá et al. 2005) |
| | Chr09:363,374 | T → G | M908L | Antimalarial multidrug resistance | Not present | (Sá et al. 2005) |
| | Chr09:364,004 | C → T | G698S | Antimalarial multidrug resistance | Reference variant | (Sá et al. 2005) |
| | Chr09:364,598 | C → T | D500N | Antimalarial multidrug resistance | Not present | (Barnadas et al. 2008) |
| | Chr09:364,557 | A → T | S513R | Resistance to Chloroquine | Reference variant | (Sá et al. 2005) |
| | Chr09:365,435 | C → A | V221L | Unknown effect | Reference variant | (Orjuela-Sánchez et al. 2009) |
| Dihydrofolate Reductase (PvDHFR) | Chr09:964,633 | C → T | A15V | Pyrimethamine resistance | Reference variant | (Ganguly et al. 2014) |
| | Chr09:964,758 | T → A | F57I | Resistance to Antifolates | Reference variant | (Imwong et al. 2003) |
| | Chr09:964,760 | C → A, C → G | F57L | Resistance to Antifolates | Reference variant | (de Pecoulas et al. 1998) |
| | Chr09:964,761 | A → C | S58R | Pyrimethamine resistance | Reference variant | (de Pecoulas et al. 1998) |
| | Chr09:964,763 | C → A, C → G | S58R | Pyrimethamine resistance | Reference variant | (de Pecoulas et al. 1998) |
| | Chr09:964,771 | C → T | T61M | Resistance to Antifolates | Reference variant | (Imwong et al. 2003) |
| | Chr09:964,884 | C → T | H99N | Unknown effect | Reference variant | (Huang et al. 2014) |
| | Chr09:964,885 | C → A | H99N | Unknown effect | Reference variant | (Huang et al. 2014) |

| | | | | | | |
|---|---|---|---|---|---|---|
| | Chr09:964,939 | G → C, G → A | S117N | Pyrimethamine resistance | Reference variant | (de Pecoulas et al. 1998) |
| | Chr09:965,106 | A → C | I173L | Resistance to Antifolates | Reference variant | (Leartsakulpanich et al. 2002) |
| **Dihydropteroate Synthase (PvDHPS)** | Chr14:1,257,064 | G → A | A647V | Unknown effect | Reference variant | (Menegon et al. 2006) |
| | Chr14:1,257,254 | T → C | I584V | Resistance to Sulfadoxine | Reference variant | (Hawkins et al. 2009) |
| | Chr14:1,257,346 | G → C | A553G | Loss of affinity for Sulfadoxine | Not present | (Korsinczky et al. 2004) |
| | Chr14:1,257,856 | G → C | A383G | Loss of affinity for Sulfadoxine | Reference variant | (Korsinczky et al. 2004) |
| | Chr14:1,257,859 | G → C | S382C | Resistance to Sulfadoxine | Reference variant | (Hawkins et al. 2009) |
| | Chr14:1,257,860 | A → C | S382A | Resistance to Sulfadoxine | Reference variant | (Hawkins et al. 2009) |
| | Chr14:1,258,389 | C → T | M 205I | Unknown effect | **Mutated variant** | (Hawkins et al. 2009) |

**Supplementary Table 9** (see extended excel file)
List of nonsynonymous SNPs, insertions, and deletions in 22 genes identified as, or being associated with, genes having high *Fst*. The reference allele frequency/alternative allele frequency is indicated (ref/alt), while "ref in all" indicates no observed SNPs at this location in this population.

**Supplementary Table 10** (see extended excel file)
Samples used in the Population Genomics Dataset.
